# Stable but turbulent: the two faces of the germline-restricted chromosome of passerine birds

**DOI:** 10.64898/2026.02.16.706238

**Authors:** S. A. Schlebusch, Z. Halenková, H. Moreno, J. Rídl, O. Kauzál, A. Suh, K. Janko, D. Dedukh, J. Pačes, T. Albrecht, R. Reifová

**Affiliations:** Department of Zoology, Faculty of Science, Charles University, 12800 Prague, Czech Republic; Laboratory of Genomics and Bioinformatics, Institute of Molecular Genetics, Czech Academy of Sciences, Prague, Czech Republic; Institute of Vertebrate Biology, Czech Academy of Sciences, Studenec, Czech Republic; Centre for Molecular Biodiversity Research, Leibniz Institute for the Analysis of Biodiversity Change, Zoologisches Forschungsmuseum A. Koenig, Adenauerallee 160, Bonn, 53113, Germany; Bonn Institute for Organismic Biology (BIOB) – Animal Biodiversity, University of Bonn, An der Immenburg 1, 53121 Bonn, Germany; Laboratory of Non-Mendelian Evolution, Institute of Animal Physiology and Genetics, The Czech Academy of Sciences, 27721 Liběchov, Czech Republic; Department of Biology and Ecology, Faculty of Science, University of Ostrava, 70103 Ostrava, Czech Republic; Institute of Molecular Genetics of the Czech Academy of Sciences, 14220 Prague, Czech Republic

## Abstract

Germline-restricted chromosomes (GRCs) are essential, supernumerary chromosomes that undergo programmed elimination in somatic cells and are only retained in the germline. Despite their recurrent emergence across animals, their genetic composition, function and evolution remain poorly understood. Here we present the most complete and contiguous GRC assemblies, including one nearly telomere-to-telomere GRC assembly, from four closely related passerine bird species, providing an unprecedented insight into the GRC’s composition and its evolution over short evolutionary timescales. We show that the passerine GRC is highly enriched in repetitive sequences, with massive, species-specific satellite expansions resulting in enormous differences in GRC size among species. Among mostly recently added sequences, we found only two ancestral genes dating back to the presumed GRC origin, offering clues to its essential function. Importantly, we demonstrate that the GRC undergoes extensive fine-scale within-chromosome rearrangements and copy number changes resulting in little collinearity between species. Our findings indicate that programmed DNA elimination has profoundly changed the GRC’s evolution by altering the selection pressures and mutational mechanisms it is exposed to. This makes the GRC an extraordinarily dynamic element in an otherwise stable avian karyotype, retaining core functions while diversifying rapidly, with important implications for germline biology, adaptive evolution and speciation.

## Introduction

Programmed DNA elimination (PDE) is an important developmental process, repeatedly evolved across multicellular organisms, in which specific genomic regions are removed from certain cell lineages (Drotos et al., 2022; Wang & Davis, 2014). The scale and architecture of elimination vary widely across taxa, ranging from excision of numerous short, dispersed fragments to the removal of entire chromosomes (Dedukh & Krasikova, 2021). PDE most commonly occurs in somatic cells, preserving the full genome only in the germline. However, germ cell-specific, usually sex-biased, elimination systems also exist. Germline-restricted chromosomes (GRCs) exemplify a notable form of PDE of entire chromosomes. They are retained exclusively in the germline and eliminated from somatic tissues. GRCs have evolved independently several times in both vertebrates and invertebrates, in some cases persisting for tens of millions of years (Hodson & Ross, 2021; Smith et al., 2021; Timoshevskiy et al., 2017; Torgasheva et al., 2019). Yet the mechanisms of their origin, reasons for their long-term maintenance and evolutionary impacts remain largely unresolved.

The passerine birds, comprising approximately 6,700 species, form one of the most species-rich clades known to carry the GRC. In all species, a single GRC is eliminated during early embryogenesis from the somatic cells (Dedukh et al., 2025) and subsequently also from male germ cells during spermatogenesis (Pigozzi & Solari, 2005; Borodin et al., 2022). Elimination proceeds via the formation of a micronucleus, where the GRC undergoes extensive fragmentation (Dedukh et al., 2025; Schoenmakers et al., 2010). Although GRC elimination seems to be highly efficient in somatic cells, it can fail in a small fraction of spermatocytes in some male individuals, enabling rare, but detectable, paternal transmission besides predominant maternal inheritance (Pei et al., 2022).

The passerine GRC was discovered more than two decades ago in zebra finch (*Taeniopygia guttata*) (Pigozzi & Solari, 1998) and its presence was recently confirmed in other passerine species, both from the oscine (Torgasheva et al., 2019) and suboscine (Ruiz-Ruano et al., 2025) passerine suborders. It is thus thought to have originated approximately 50 Mya in the common ancestor of all passerines, with the possible exception of the small basal lineage, Acanthisittidae, which has not been studied yet (Schlebusch et al., 2023; Ruiz-Ruano et al., 2025). Since all passerine species investigated so far possess the GRC, it is believed to be essential for these birds, although its functions remain unknown. The GRC’s genetic content appears to be largely, if not entirely, composed of sequences coming from regular chromosomes (i.e. the autosomes and sex chromosomes) also called A chromosomes, which have been successively copied onto the GRC during passerine evolution (Kinsella et al., 2019; Mueller et al., 2023; Schlebusch et al., 2023). Although most genes identified on the GRC seem to be nonfunctional truncated pseudogenes (Schlebusch et al., 2023), several important genes with complete coding regions and expression in the germline have been identified on the GRC (Kinsella et al., 2019; Mueller et al., 2023; Schlebusch et al., 2023; Vontzou et al., 2025). Some of these genes have diverged substantially from their A-chromosomal paralogs, potentially acquiring novel germline-specific functions (Schlebusch et al., 2023). In rare cases, an A-chromosomal paralog has been lost after being duplicated on the GRC (Ruiz-Ruano et al., 2025). These genes could underpin the indispensability of the passerine GRC, yet their specific roles and functional divergence from A-chromosomal counterparts remain unclear.

Despite its apparent indispensability, the passerine GRC exhibits remarkable variation in both size (Torgasheva et al., 2019) and genetic content (Schlebusch et al., 2023; Ruiz-Ruano et al., 2025). The size of the GRC ranges from one of the largest chromosomes (referred to as macro GRCs) to one of the smallest chromosomes (micro GRCs) in the germline karyotype. This variability is evident even over short evolutionary time scales, such as among closely related estrildid finches of the *Lonchura* genus (Sotelo-Muñoz et al., 2022). Genetic composition of the GRC can also differ substantially, even among species with similarly sized GRCs. For example in closely related nightingale species, one-third to one-half of the GRC sequence is species-specific (Schlebusch et al., 2023), and only a minute fraction of the GRC is shared between two main passerine suborders, oscines and suboscines (Ruiz-Ruano et al., 2025). This extreme variability contrasts sharply with the highly conserved karyotypes of birds, characterized by stable chromosome numbers and sizes, and extensive sequence synteny even across deeply diverged lineages (Zhang et al., 2000; Kapusta & Suh, 2016). The mechanisms driving this exceptionally fast evolutionary dynamics as well as its potential evolutionary consequences remain unclear, underscoring the need for comparative analyses, especially among closely related species.

It has been proposed that the GRC may have originated as a selfish B chromosome which evolved somatic elimination to be less harmful to its host and acquired important functions preventing its loss from the genome (Johnson Pokorná & Reifová, 2021). It has been thus assumed that the ancient genes responsible for the GRC’s initial origin and ongoing retention would be shared among passerine species. However, a recent comparative study of the GRC genetic content across major passerine lineages surprisingly revealed no shared GRC genes among all the analysed species (Ruiz-Ruano et al., 2025). This may either reflect rapid GRC gene content turnover and the existence of different important genes in different passerine lineages (Schlebusch et al., 2023; Ruiz-Ruano et al., 2025) or, alternatively, limitations of the existing GRC assemblies, which are mostly incomplete and fragmented. The poor quality of existing GRC assemblies is mostly the result of the A-chromosomal origin of the GRC and the fact that the GRC is only found in a small fraction of cells in gonadal tissue, which makes the GRC difficult to sequence and assemble. This is especially true for short read data, which most of the assemblies are based on. High-quality, contiguous GRC assemblies are critical to resolving the full GRC’s genetic content, including repetitive sequences, which are largely missing from the current GRC assemblies.

To characterize the genetic composition of the passerine GRC, explore its evolutionary dynamics, and identify important functional genes, we sequenced and assembled the GRCs of four estrildid finches of the genus *Lonchura*. Two of these species (*L. punctulata* and *L. malacca*) carry micro GRCs, whereas the other two species (*L. striata var. domestica* (hereafter *L. domestica*) and *L. castaneothorax*) possess macro GRCs (Sotelo-Muñoz et al., 2022). These species diverged approximately within the last 4 million years (Kumar et al., 2017), and their phylogeny suggests that at least two independent contractions or expansions in GRC size occurred within this time (Figure 1). This system thus offers a unique opportunity to uncover the mechanisms shaping the evolution of the essential, yet rapidly evolving GRC.

**Figure 1:**
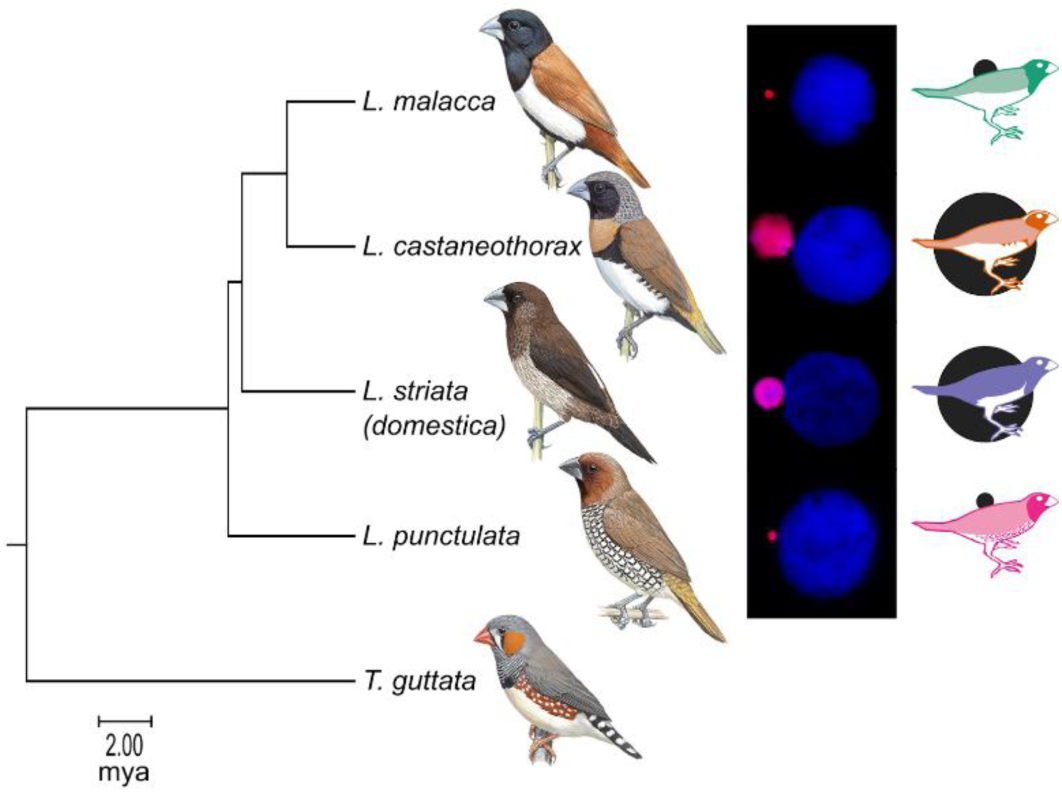
Phylogeny of the four *Lonchura* species with zebra finch (*Taeniopygia guttata*) as an outgroup. The *Lonchura* species are accompanied by photos of their GRCs being excluded as a micronucleus (stained by anti-H3S10p antibody, in pink) from nuclei of male germ cells (stained by DAPI, in blue) (taken from (Sotelo-Muñoz et al., 2022)). The size of the micronuclei reflects the size of the GRC. The outgroup, *T. guttata*, is known to have a macro GRC (Pigozzi & Solari, 1998). The simplified illustrations of each species with indicated GRC size in the background (on the right) is used to represent the species in the following figures. The divergence times used were estimated with TimeTree (Kumar et al., 2017). Scale bar shows million years of divergence. Detailed illustrations courtesy of Cornell Lab of Ornithology, Hilary Burn from Birds of the World (https://doi.org/10.2173/bow)

Our assemblies, particularly those of the micro GRCs, represent the most contiguous and complete GRC assemblies reported to date. Notably, the *L. malacca* micro GRC approaches the first telomere-to-telomere assembly of a GRC, not only in passerine birds but generally in animals. Using these high-quality assemblies, we explored the evolutionary dynamics of the GRC over short evolutionary distances, clarifying both conserved features and lineage-specific shifts in gene and repeat content. The exceptional assembly quality further enabled the first cross-species analysis of whole-chromosome GRC synteny and structural changes, revealing extensive within-chromosome rearrangements and copy-number variation despite general conservation of genetic content. Together, our findings establish the GRC as a distinctive component of the passerine genome, characterized by highly dynamic repeat as well as gene content, and pervasive intra-chromosomal restructuring, yet maintaining its core essential genes mostly regulating gene expression at the translation level during early embryogenesis.

## Results

### GRC assemblies

In order to assemble the GRC of the four *Lonchura* species, PacBio HiFi sequencing of testis and kidney tissue was done for each species. The testis data was then assembled such that similar sequences were not merged like they normally would be during the assembly process. This was done to prevent chimeric GRC/A-chromosomal contigs being created from GRC sequences that are highly similar to their A-chromosomal counterparts. These testis assemblies were all about 2.3 Gbp, consistent with maternal and paternal haplotypes being distinguished in the assembly process (for reference, *T. guttata*’s genome size is 1.14 Gbp (Formenti et al., 2025)). Any contigs which had a normalised depth 1.5x higher in testis than in kidney were selected to form an initial set of GRC contigs. These contigs were then manually inspected and further refined using additional information such as Omni-C connections and the proportion of kmers unique to the testis (see Materials and Methods for more details). The final GRC assembly lengths (see Table 1) were consistent with our expectation of macro and micro GRC sizes based on previous cytogenetic results (Sotelo-Muñoz et al., 2022).

**Table 1:** Final assembly statistics of the GRC for each *Lonchura* species. Testis and kidney read depth are reported relative to the average depth that the respective dataset had across the A chromosomes.

|  | *L. malacca* | *L. punctulata* | *L. castaneothorax* | *L. domestica* |
| --- | --- | --- | --- | --- |
| GRC length | 8.2 Mbp | 4.4 Mbp | 150 Mbp | 130 Mbp |
| No. of contigs | 8 | 12 | 499 | 584 |
| N50 | 2.5 Mbp | 810 kbp | 560 kbp | 540 kbp |
| N90 | 390 kbp | 180 kbp | 140 kbp | 110 kbp |
| Testis depth | 71% | 46% | 32% | 48% |
| Kidney depth | 0.1% | 1.4% | 0.0% | 3.2% |

While the two macro GRCs consisted of hundreds of contigs, the two micro GRCs were substantially more contiguous, with *L. malacca* having 8 contigs and *L. punctulata* having 12 contigs (Table 1, Figure 2 and Supplementary Figures 1-5). With the help of Omni-C data with 150-bp and 300-bp reads we were able to further scaffold these contigs. In the case of *L. malacca*, this resulted in a chromosome-level assembly with telomere repeats found at both ends (Figure 2).

**Figure 2:**
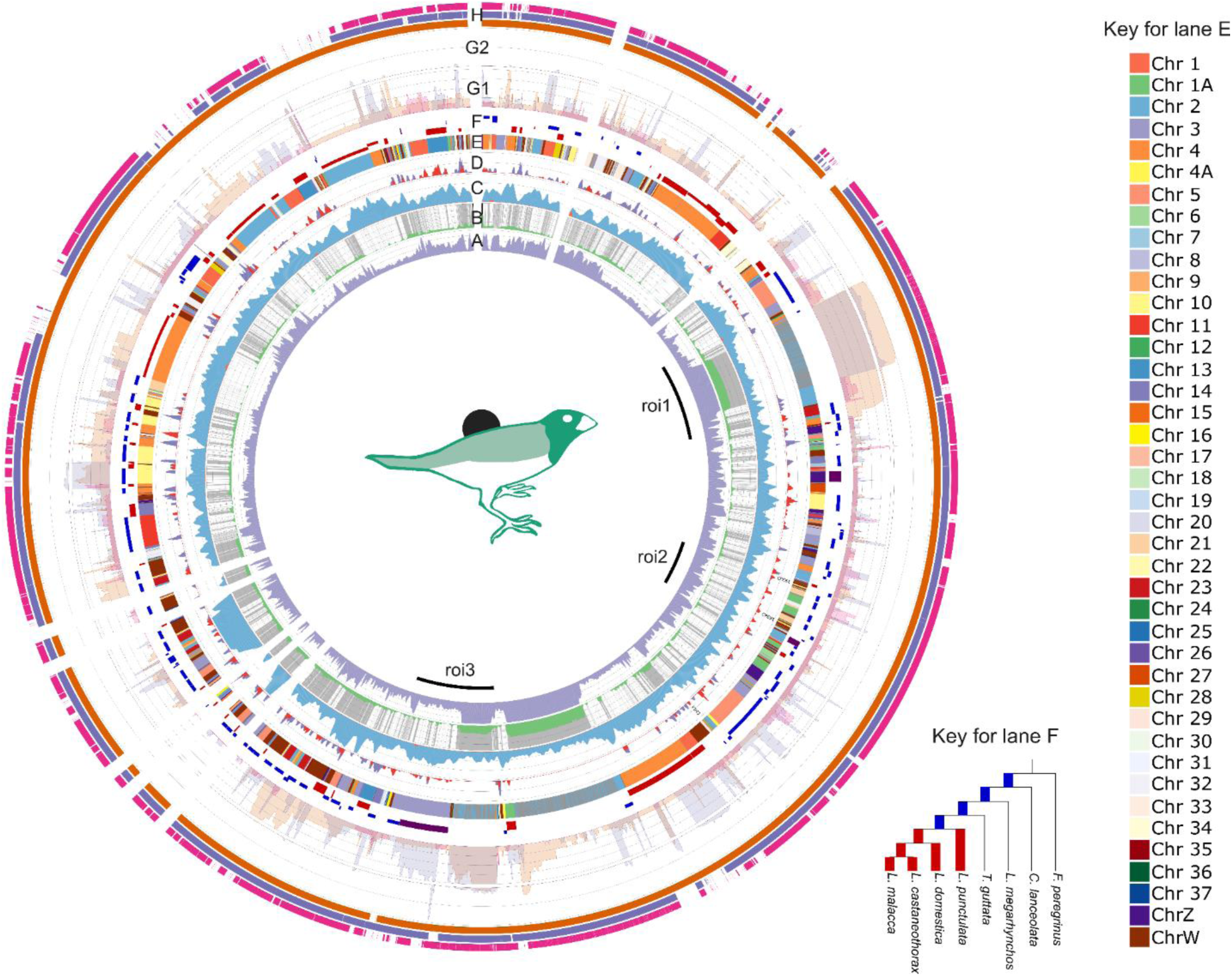
The *L. malacca* GRC assembly and its characteristics. A) Proportion of 35-bp kmers which were unique to the testis dataset. B) Average number of times (log scale) 35-bp kmers were found on the whole GRC (green) as well as whether a region was annotated as repetitive (grey). Telomeres are shown in black (and flank the letter B). C) Normalised coverage of the GRC contigs from the testis dataset (blue) and kidney dataset (orange) as a proportion of their coverage across the A chromosomes. D) Gene density for all identified genes (purple) as well as only functional genes (red). E) A chromosome that the GRC sequence aligned to (see Key for lane E). F) Estimated age of the GRC region. The young sequence (red) originated within the diversification of the *Lonchura* species (i.e. within the last 4 million years), while the older sequence (blue) predates the diversification. If there was ambiguity about whether a sequence diverged from the A chromosomes before or after the Lonchura diversification, it is shown in purple. The further away from the center of the Circos plot, the older the region (see Key for lane F). G) Alignment of the other *Lonchura* species’ GRCs to *L. malacca*. This is shown on a linear scale (G1) for less than or equal to 8 alignments, after which it is shown on a log scale (G2). H) Indicates whether the *L. malacca* GRC aligned to *L. castaneothorax* (orange), *L. domestica* (purple), and *L. punctulata* (pink). Lanes A, B, C, and D used a sliding window of 10 kbp and a step size of 2 kbp (minimum size 5 kbp). Lane F and G used a sliding window of 2.5 kbp and a step size of 500 bp (minimum size 1 kbp). Three regions of interest (roi) are shown inside the plot showing the location of two notable repeat structures (roi1 and roi3) and the location of the oldest identified region (roi2).

GRC sequences were found to have a consistent set of characteristics distinct from the A chromosomes. They often have low depth in the kidney dataset (1.0% of the depth of the average haplotype resolved A chromosome), while the testis dataset had an average depth of 49%, representing a substantially larger fold difference than the initial selection criteria. The contigs also consistently align to multiple A chromosomes with few long contiguous alignments. Finally, each macro GRC had its own abundant repeat present on many of the contigs (see Supplementary Figure 1 and 2).

### Characterization of the GRC repeat content

The GRCs of all four species contained a higher proportion of repeats than the A chromosomes (Figure 3). Repeats made up 25-27% of A-chromosomal sequence in the four *Lonchura* species. In contrast, 47-93% of the sequence was repetitive on the GRC. Consistent with their size, the macro GRCs had a higher proportion of repeats (93% in *L. castaneothorax*, 86% in *L. domestica*) than micro GRCs (53% in *L. malacca*, and 47% in *L. punctulata*).

**Figure 3:**
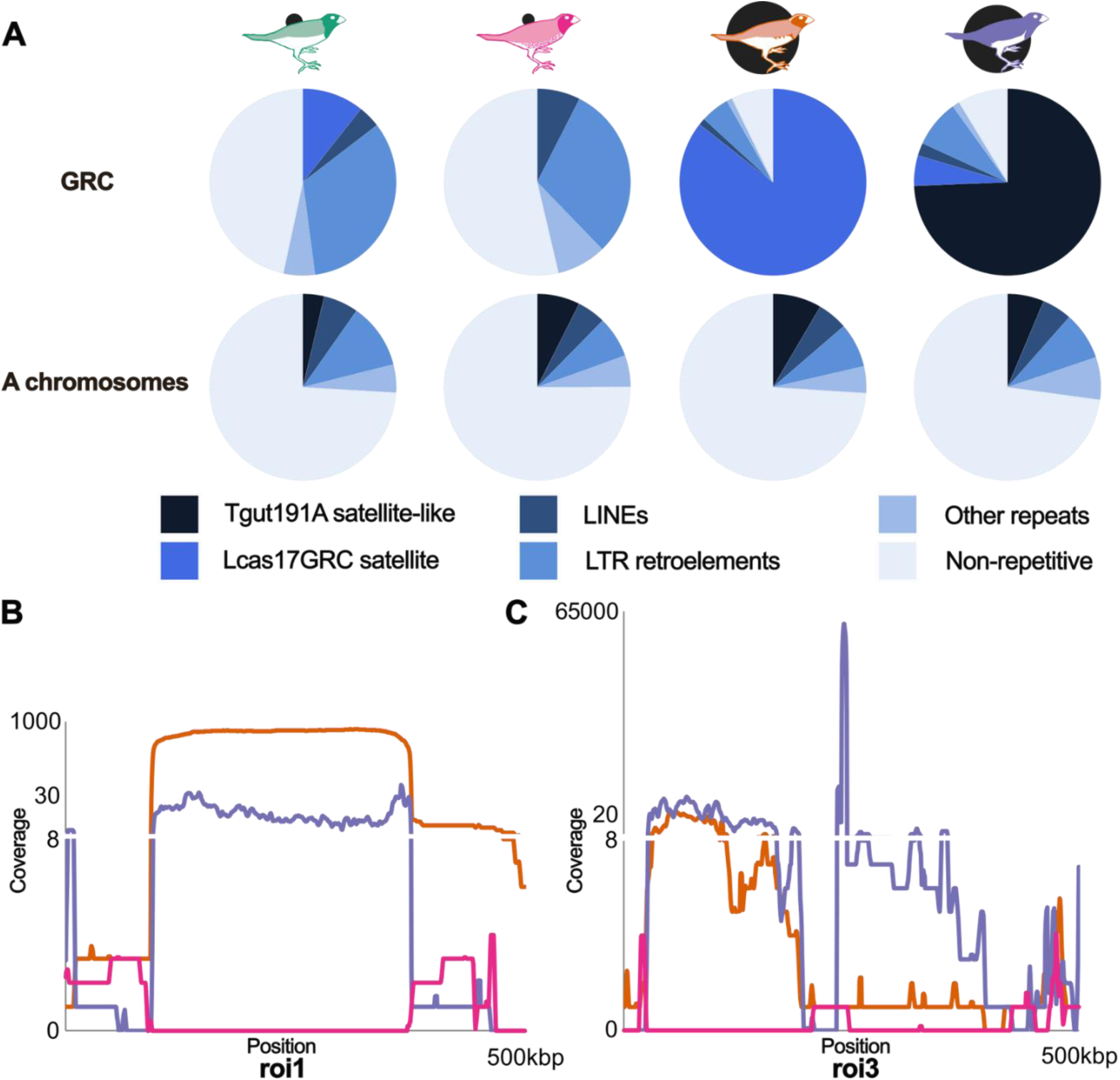
The GRC repeat content in the four *Lonchura* species. A) The proportion of each GRC assembly and A-chromosomal assembly made up of non-repetitive and repetitive DNA, including satellites, LINEs, LTR retroelements and other repeats. B) The number of times *L. punctulata*, *L. castaneothorax* and *L. domestica* aligned to roi1 (see Figure 2), which contained Lcas17GRC. C) The number of times the three *Lonchura* species aligned to roi3 (see Figure 2), which contained the Tgut191A-like satellite. Note that B and C have 1 to 8 copies on a linear scale before a break signifies a switch to a log scale in both cases.

The two macro GRCs were found to be predominantly made of satellite repeats. Each macro GRC had a single repetitive sequence that made up the vast majority of its length. Despite the two GRCs’ similar size and relatedness, the relevant repeat sequences were unrelated. *L. domestica* had a tandem repeat of 191-bp motif which was found over 520 000 times with a cumulative length of 99 Mbp (74% of the assembled GRC sequence). This sequence was reasonably well conserved, with an average fraction of matching nucleotide positions between adjacent copies of ∼86%, and an average indel rate of ∼2%. In comparison, *L. castaneothorax* had a relatively degraded 17-bp tandem repeat that covers 131 Mbp (86% of the GRC length), with an average conservation of 79% and an indel rate of 5% between adjacent copies.

Interestingly, the 191-bp repeat on the *L. domestica* GRC shared a high sequence identity with Tgut191A (Supplementary Figure 6), a centromere-associated tandem repeat from *T. guttata* found throughout the genome (Takki et al., 2022; Formenti et al., 2025). This repeat is present on the other three GRCs, but in much smaller numbers (1-7 kbp total in each GRC) (see roi3 region in Figure 2 and Figure 3 C for examples).

The *L. castaneothorax* repeat sequence was also found on the GRC of the other species except for *L. punctulata* (roi1 region in Figure 2 and Figure 3 B). Several long degenerate repeats originating from chromosome 2 were present on the *L. malacca* GRC and made up the major repeat structures visible on the chromosome (Supplementary Figure 7). All three other species’ GRCs aligned to these regions to some degree. However, *L. punctulata’s* GRC only aligned to the flanking region originating from chromosome 2, while the repetitive region was absent. *L. domestica* GRC sequence aligned to most of the relevant regions, with the repetitive sequence duplicated several times. In *L. castaneothorax*, however, one of those repeat regions (280 kbp long) was duplicated 600-700 times in roughly equal proportions (see Figure 3 C), possibly suggesting that it was duplicated as a whole and that the repeats are hierarchically structured, with the 17-bp repeat nested inside the 280-kb repeat. We called this satellite Lcas17GRC.

In addition to these extensive satellite expansions, the GRCs of all four species showed relatively higher content of LTR retroelements compared to the A chromosomes. LTR retroelements made up approximately one third of the GRC size when excluding the massive satellite array structures in *L. castaneothorax* and *L. domestica* (33% in *L. malacca*, 35% in *L. castaneothorax*, 32% in *L. domestica* and 29% in *L. punctulata*), compared to 7-11% in the A chromosomes, making a significant enrichment (p = 1.7e-4, paired t-test). There was also a small proportion of LINE elements and other repeats on the GRC (Figure 3), but the proportion of LINE elements did not significantly differ between the GRC and A chromosomes (p > 0.05, paired t-test).

### Characterization of the GRC gene content

Genes were annotated on the GRCs of the four *Lonchura* species using aligned *T. guttata* proteins. Overall, we identified 236 distinct genes on the GRCs of the four *Lonchura* species (157 in *L. domestica*, 142 in *L. castaneothorax*, 89 in *L. malacca*, and 67 in *L. punctulata*). Many of these genes (42%) were found to be duplicated on the GRC, with two or more copies present (Figure 4 A). In some cases, the number of gene copies reached several tens or hundreds (Supplementary Table 1). Despite the recent divergence of the *Lonchura* species, gene content differed considerably between the GRCs. Of the 236 distinct genes identified, only 54 were found on all four GRCs (see Supplementary Table 1). A relatively large number of genes on the GRC were species-specific, i.e. found only in one of the four species. There were more species-specific genes on macro GRCs (82 in *L. domestica* and 51 in *L. castaneothorax*) than in micro GRCs (3 in *L. malacca* and 6 in *L. punctulata*). The shared genes were disproportionately more duplicated (69% of genes duplicated) compared to all GRC genes combined (42% of genes duplicated). Species-specific genes were duplicated only 30% of the time.

**Figure 4:**
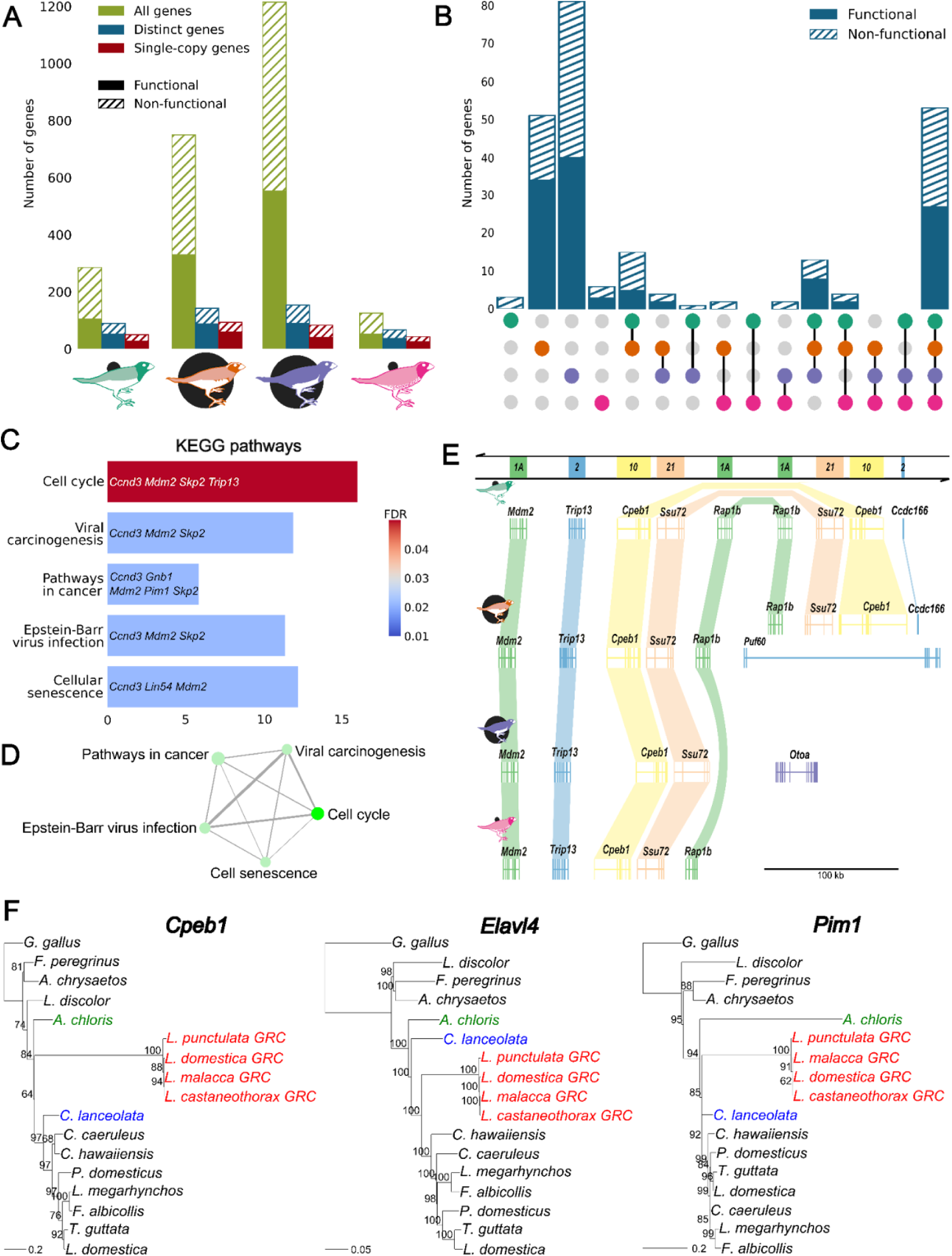
GRC gene content. A) The total number of genes (including gene duplicates) per species and whether they were classified as functional or non-functional. Distinct genes are classified as functional if they have at least one functional copy. B) Upset plot of the number of distinct genes between all possible subsets of the four GRCs. Genes are classified as functional when there is at least one functional copy in all species of the subset. C) KEGG pathways enrichment analysis of functional GRC genes shared between all four species. Bar lengths represent the fold enrichment values, with the genes from which the enrichment was based being shown on the bar. Color scale indicates the false discovery rate (FDR). Only significantly enriched KEGG pathways are shown. D) Network of enriched KEGG pathways. Each node represents an enriched pathway. Two pathways are connected if they share 20% or more genes. The size of the nodes indicates the number of genes in the corresponding pathway, while the brightness indicates the statistical significance. E) Gene content around *Cpeb1*, one of the oldest regions in each species. Genes are shown as exons (coloured boxes) and introns (connecting lines) coloured according to their A-chromosomal origin (which is also illustrated by the bar above the figure), with synteny illustrated by connections between homologs. LOC genes are not shown. F) Maximum likelihood phylogenetic trees for the only three genes with an estimated GRC acquisition before the radiation of oscines. GRC genes are highlighted in red, the suboscine sequence in blue, and the *Acanthisitta chloris* sequence in green. Bootstrap support values are indicated at the nodes. *Elavl4* is based on the complete gene sequences (including introns) to solve ambiguity in the divergence timing.

Each annotated gene was then classified as functional or non-functional. We considered a gene to be functional if its coding sequence covered more than 90% of the length of the A-chromosomal paralog and did not contain any premature stop codons or frame-shift mutations within the first 90% of the coding region. We considered all other cases as non-functional. The majority of all GRC genes (56%) were non-functional according to these criteria. This pseudogenization was usually caused by severe shortening of the coding sequence (see Supplementary Figure 8 for breakdown) and mostly occurred in multicopy genes. Overall, 91% of non-functional genes were from multicopy GRC genes, while only 9% were single-copy genes. Overall, 55-63% of the distinct GRC genes had at least one functional copy in a species (see Figure 4 A).

Of 54 shared genes among all four *Lonchura* species, 28 (52%) had at least one functional copy in all four species, and 13 (24%) had no functional copy in any species (Figure 4 B). Of the 142 species-specific genes, 54% had at least one functional copy. Interestingly, there is generally a consensus among the GRCs about whether a shared gene is functional or not. Of genes shared by at least two species, 31% are pseudogenized in all those species and 47% are functional in all those species (i.e. have at least one functional copy). This means that whether a shared gene is functional or not only differs in 22% of genes, even though the vast majority of genes are recent additions to the GRC, that were duplicated on the GRC during the *Lonchura* radiation (see below), which limits their potential shared evolutionary history in common ancestors.

GRC genes had fewer introns than their A-chromosomal reference, with a median difference of 2 introns (Mann–Whitney U test, p = 1.8e-3). This was true after taking into account the general truncation of genes observed on the GRC. However, this pattern was not observed when only looking at functional genes (Mann–Whitney U test, p > 0.05) or the longest version of a gene (Mann–Whitney U test, p > 0.05). Introns were also found to be consistently shorter on the GRC compared to A chromosomes, with introns in general having been 290 bp shorter across all GRC genes (Mann–Whitney U test, p = 1.2e-10), 339 bp shorter in functional genes (Mann–Whitney U test, p = 1.5e-14) and 256 bp shorter in the longest version of a gene (Mann–Whitney U test, p = 3.4e-9), consistent with findings from Mueller et al (2023) (see Supplementary Figure 9 A and B). However, despite the average intron size reduction on the GRCs, there was high variation at the gene level, with the three oldest genes identified on the GRC (*Cpeb1*, *Elavl4*, *Pim1*) (see below) having mostly longer, not shorter, introns than their A-chromosomal paralogs (see Supplementary Figure 9 C, D and E).

The functional GRC genes identified across all four *Lonchura* species or in individual *Lonchura* species did not show any significant enrichment for GO terms or KEGG pathways. When limited to only the 27 shared functional genes, there was still no GO term enriched, but there was significant enrichment for the following KEGG pathways: cell cycle, cellular senescence, viral carcinogenesis, Epstein-Barr virus infection and pathways in cancer (Figure 4 C and D). These pathways were enriched based on the following genes, *Ccnd3*, *Gnb1*, *Lin54*, *Mdm2*, *Skp2*, *Pim1* and *Trip13*. Two of these seven genes (*Mdm2* and *Trip13*) were clustered adjacent to each other (see Figure 4 E) along with the mRNA-binding protein *Cpeb1*, and the RNA Polymerase II phosphatase *Ssu72* in all four species.

### Age of GRC sequences

Across each GRC, we estimated the evolutionary period in which each sequence was duplicated onto the GRC from A chromosomes by comparing its evolutionary relationship to A-chromosomal sequences from multiple species covering the Passeriformes order. These included the 4 *Lonchura* species, *T. guttata* and *Luscinia megarhynchos* representing an oscine order, *Chiroxiphia lanceolata* representing a suboscine order, and *Falco peregrinus* as a non-passerine outgroup. The divergence of the GRC sequence in this genealogy indicated the period when the sequence was copied from the A chromosomes to the GRC (see Figure 2).

Consistently with the previous work (Kinsella et al. 2019, Schlebusch et al. 2023), we found that a large proportion of each GRC was duplicated from A chromosomes recently in evolutionary time. For example, in *L. malacca*, where we were able to estimate the age of divergence of 4.8 Mbp (59% of the total GRC length and disproportionately non-repetitive), approximately 44-49% of this sequence was estimated to have originated within the last 4 million years during the radiation of the *Lonchura* genus. In comparison, only two small regions with a total length of 26 kb (0.5% of DNA with an age estimate) were estimated to be older than the oscine/suboscine divergence, i.e. older than 44 million years (Figure 2). These regions corresponded exactly with the *Cpeb1* gene, which had been duplicated in this species (Figure 4 E).

*L. punctulata* had the least new sequences, with only 27-32% of the 2.8 Mbp with an age estimate originating within the last 4 million years (Supplementary Figure 5). In comparison, the two macro GRCs had the largest fraction of new sequences with 67-70% of 15 Mbp (the amount that had an age estimate assigned to it) in the case of *L. castaneothorax* and 81% of 18 Mbp for *L. domestica* (Supplementary Figures 3, 4). Like in *L. malacca*, the proportion of old sequences (from before the divergence of suboscines and oscines) on the GRC of the other three species was consistently small, between 11 kb and 34 kb, and corresponded with *Cpeb1* gene, which was present in a single copy in *L. domestica* and *L. punctulata*, and two copies in *L. castaneothorax* (Figure 4 E). Across all four GRCs, newer sequences often made longer and more continuous blocks than older sequences, which were smaller and often had other sequences of various ages interspersed with them (Figure 2).

Focusing on GRC coding sequences, we further estimated the age of individual genes shared by the four species and functional in at least three of them, by creating their genealogies with A-chromosomal paralogs from 13 avian species. This gene-based approach can also reveal old GRC genes not identified on the coarser whole chromosomal scale. Of the 32 genealogies created, only two supported a duplication to the GRC before the suboscine and oscine split. These corresponded to *Cpeb1 and Pim1* genes. For another one, it occurred before the oscine radiation (*Elavl4*) (see Figure 4 F). Six other genes (*Lin54*, *Mdm2*, *Pim3*, *Rabac1*, *Skp2* and *Ugdh*) were duplicated onto the GRC before the divergence of *Lonchura* lineage from *T. guttata*, while 15 genes were recent acquisitions that originated after the split from *T. guttata (*see Supplementary Figure 10). The remaining 8 genes (*Erfl*, *Gabarap*, *Gnb1*, *Phlda1*, *Rap1b*, *Ssu72*, *Suv39h1*, *Trip13*) are relatively recent acquisitions but their genealogies were not able to clearly estimate the time of the duplication event.

### Structural variation on the GRC between species

Despite the large discrepancy in size and the difference in gene content, the overall genetic content of each species’ GRC was surprisingly similar, with 96% of the *L. castaneothorax* GRC and 93% of the *L. domestica* GRC aligning to the *L. malacca* GRC. Consistent with their phylogeny (Figure 1, Stryjewski & Sorenson, 2017), *L. malacca* aligned best to *L. castaneothorax* (99%) and worst to *L. punctulata* (74%), while *L. punctulata* aligned equally well to all the other GRCs (Figure 5 B). The two macro GRCs alignment rates were not what would be predicted by the phylogeny, but this is consistent with their sequence mostly being repeats that either show or lack the relevant homologous sequence in the other species (see Figure 5 B).

**Figure 5:**
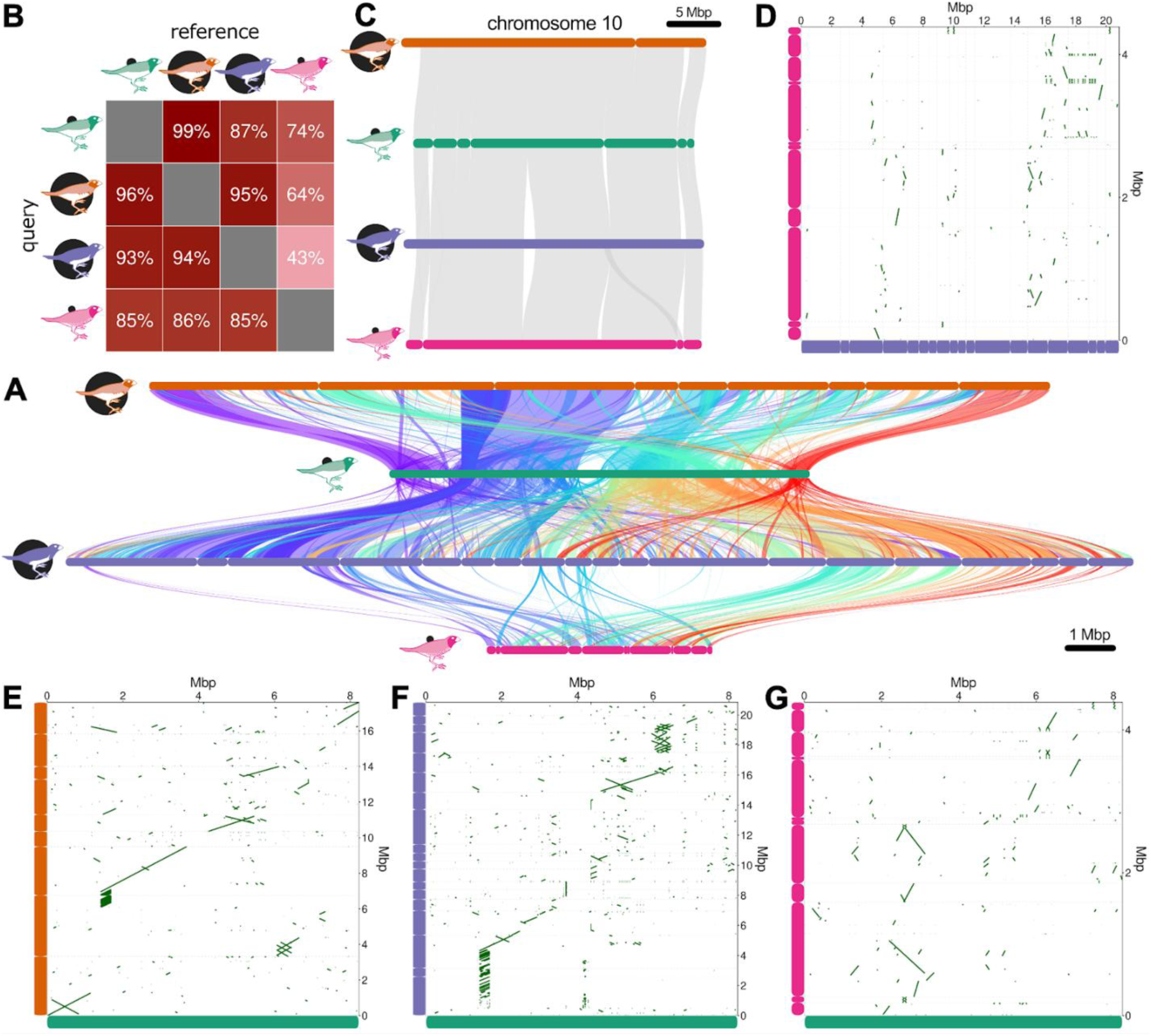
Shared synteny between the *Lonchura* GRCs. A) Collinearity plot of GRC sequences of *L. castaneothorax*, *L. malacca*, *L. domestica* and *L. punctulata* (top to bottom). Due to the size and repetitive nature of *L. domestica* and *L. castaneothorax*, only contigs that were larger than 500 kbp and had less repetitive 35-bp kmers than the contigs that contained *Cpeb1* were selected (see Supplementary Figures 1-4 for visualisations). Contigs of *L. castaneothorax, L. domestica,* and *L. punctulata* were ordered and oriented according to the alignment with the chromosome-level GRC assembly of *L. malacca*, thus maximizing collinearity between the species. This figure thus represents a lower estimate of the amount of rearrangements on GRC. The color of the connection is determined by the reference position from left to right. The synteny plot was generated using NGenomeSyn (He et al., 2023). B) Proportion of aligned sequence between individual GRCs. C) Synteny between the four *Lonchura* species in a selected A-chromosome (chromosome 10). A-chromosomal contigs of *L. castaneothorax*, *L. malacca* and *L. punctulata* were ordered and oriented according to the chromosome 10 of the published assembly of *L. striata* (NCBI RefSeq ID GCF_046129695). Only contigs longer than 500 kbp and aligning with *L. striata*’s chromosome 10 in at least 50% of their length were selected. D-F) Dotplots visualising the synteny between the GRC of *L. domestica* (subset) and *L. punctulata* (D) and *L. malacca* and *L. castaneothorax* (subset) (E), *L. malacca* and *L. domestica* (subset) (F), and *L. malacca* and *L. punctulata* (G). Contigs of *L. castaneothorax*, *L. domestica* and *L. punctulata* are ordered in the same way as in panel A. Dotplots were generated using shinyChromosome (Yu et al., 2020).

Given that the *L. malacca* GRC was the best GRC assembly and the other assemblies align well to it, we used it as a reference to check for synteny and presence of structural variation across the four species. With the more repetitive contigs excluded from the two macro GRCs, relatively well conserved synteny between *L. malacca*’s GRC and the closely related *L. castaneothorax* GRC was visible (Figure 6 A and E). The order and orientation of sequences was, however, still much more disorganized than in A chromosomes (Figure 5 C). The synteny of *L. malacca’s* GRC was greatly reduced with the more distantly related *L. domestica* (Figure 5 A and F) and almost non-existent with *L. punctulata* (Figure 5 D and G and Supplementary Figure 11).

## Discussion

A growing body of evidence suggests that somatic programmed DNA elimination, restricting parts of the genome to the germline, is a widespread phenomenon present in thousands of species across vertebrates and invertebrates (Wang & Davis, 2014), with GRCs having evolved independently in passerine birds (Kinsella et al., 2019; Torgasheva et al., 2019), lamprey and hagfish (Smith et al., 2021), and three clades of Diptera flies (Hodson & Ross, 2021). The reasons for this elimination as well as its evolutionary consequences are, however, still not well understood. Using accurate long-read sequencing, we produced high-quality assemblies of the passerine GRC in four closely related *Lonchura* species differing markedly in GRC size. In one of these species, *L. malacca,* we managed to arrange the GRC contigs into a single chromosome model with seven gaps and with telomeres found on both ends by using Omni-C data. This represents the first nearly telomere-to-telomere assembly not just of a passerine GRC, but of any GRC. These assemblies have given us an unprecedented view of the GRC’s composition and its evolution.

One of the most noticeable features of the passerine GRC’s genetic content is the massive array of tandem repeats which make up the vast majority of the two macro GRCs. Notably, the nature of the repeat is different in the two *Lonchura* species, with *L. domestica* having 191-bp satellite repeat (covering 99 Mbp) and *L. castaneothorax* a degenerate 17-bp Lcas17GRC repeat forming an array of 131 Mbp length. Interestingly, the 191-bp satellite in *L. domestica* shows high similarity to the centromere-associated Tgut191A satellite identified in *T. guttata* (Takki et al., 2022). Massive amplification of a satellite repeat has been observed also in *T. guttata,* which also possesses a macro GRC. Its sequence is unrelated to repeats found in *Lonchura* species and includes an inactivated gene (*Dph6*) duplicated hundreds of times (Kinsella et al., 2019). These findings suggest that macro GRCs can rapidly evolve from micro GRCs when a single sequence undergoes massive amplification. Mechanisms leading to such amplification are not clear, but could involve unequal crossing-over during female meiosis, where a GRC pairs and recombines with its identical copy that arise by duplication of the chromosome prior to meiosis (Borodin et al., 2022), or formation of extrachromosomal circular DNA molecules and their rolling-circle replication (Navrátilová et al., 2008). Bird chromosomes are generally conserved in their length (Poignet et al., 2021), possibly due to constraints on the genome size (Wright et al., 2014). However, these constraints do not need to be equally strong in the germline, possibly providing a permissive environment for dramatic, lineage-specific repeat accumulation and rapid GRC size evolution.

In addition to accumulation of satellite repeats on macro GRSs, the GRCs of all four *Lonchura* species were found to be enriched in LTR retroelements, which constitute about one third of the GRCs’ sequence after excluding the massive satellite arrays on the macro GRCs. Our results are consistent with a very recent study in house finch (*Haemorhous mexicanus*), common rosefinch (*Carpodacus erythrinus*) and blue tit (*Cyanistes caeruleus)* showing that the GRC may be a reservoir of intact endogenous retroviruses (ERVs), which are typically removed by ectopic recombination between LTRs on A chromosomes (Fang & Edwards, 2025). The reasons for the accumulation of LTR retroelements on the GRC are unclear, but it is possible that these selfish genetic elements hijack the GRC to propagate themselves through the germline, while limiting their potential negative impact on the host in somatic cells. If this is true, it would be mirroring the same set of selection pressures that have been suggested to have led to the origin of the GRC itself (Johnson Pokorná & Reifová, 2021). Another possibility is that, as with satellites, reduced selection pressure on GRC size due to its presence only in the germline may allow accumulation of LTR retroelements on the GRC. We cannot, however, also rule out the possibility that individual LTR retroelement copies have some function on the GRC given that they sometimes form important components of heterochromatin or centromeres (Talbert & Henikoff, 2020).

Consistent with what has been previously reported for two closely related *Luscinia* species (Schlebusch et al., 2023), the gene content of the GRC in the four *Lonchura* species is remarkably different, with a relatively large number of genes being species-specific (60%). Most of the genes identified on the GRC show high copy-number variation, and many genes are truncated, have premature stop codons, or have frameshift mutations, resulting in a high abundance of pseudogenes on the GRC (56% of all genes). Of 236 distinct genes identified across all four GRCs, only 28 were shared among all four species with at least one functional copy in each species. Interestingly, these genes are significantly enriched in functions related to cell cycle, carcinogenesis and cellular senescence. This is consistent with the idea that restriction of the GRC to germ cells may allow for the accumulation of genes with negative effects in the soma and beneficial effects in the germline, such as germ cell cancer genes (Vontzou et al., 2023). Indeed, among the genes responsible for the functional enrichment paralogs of several well known oncogenes were found (e.g. *Ccnd3*, *Mdm2, Skp2 or Pim1*). These genes are known for their function as cell cycle regulators and their overexpression promotes cell division and cancer development (Croce, 2008). Another notable gene related to cell cycle control on the GRC (located right next to *Cpeb1* on the GRC of all four *Lonchura* species) is a paralog of the *Trip13* gene, which not only functions as an important cell-cycle checkpoint controlling kinetochore assembly and chromosome segregation during mitosis, but also regulates dsDNA break repair and inactivation of unsynapsed chromosomes in meiosis (Lu et al., 2019). This gene could potentially be involved in GRC elimination from somatic and male germ cells, which, in the case of somatic elimination, is known to be associated with a delay in the attachment of the GRC to the spindle and its resultant missegregation in the anaphase (Dedukh et al., 2025). In addition, during male meiosis, aberrant kinetochore assembly was observed on the GRC in pachytene, possibly resulting in failure of spindle attachment and subsequent loss of the GRC (Schoenmakers et al., 2010).

The majority of the GRC in the four *Lonchura* species have been duplicated onto the GRC from A chromosomes after the divergence of *Lonchura* species from *T. guttata*, which means that they are younger than 10 million years according to the estrildid finch phylogeny (Olsson & Alström, 2020). This makes the *Lonchura* GRC very different from that of the closely related *T. guttata* and explains why it carries so many species-specific genes. Within this predominantly young GRC content, we found only three old functional genes shared among the four *Lonchura* species, which were added to the GRC before the radiation of oscines. Two of these genes (paralogs of *Cpeb1* and *Pim1* genes) have been added onto the GRC in the ancestor of suboscines and oscines, i.e. more than 44 million years ago (Oliveros et al., 2019), in the estimated period of the GRC origin. The last one (paralog of *Elavl4* gene) was added slightly later in the ancestor of all oscines, i.e. 38–44 million years ago (Oliveros et al., 2019). *Cpeb1* was recently identified as the only complete, single-copy gene shared among two closely related *Luscinia* species (Schlebusch et al., 2023) and one of the most commonly found genes on the GRC across the major lineages of passerines (Ruiz-Ruano et al., 2025). *Cpeb1* is known to play a crucial role in orchestrating protein synthesis during oocyte maturation and early embryogenesis, when transcription is suppressed and protein synthesis depends on the translation of prestored mRNAs (Rouhana et al., 2023). Interestingly, *Elavl4* has a similar function and regulates the translation of mRNAs by recognizing similar motifs on mRNAs’ 3’-UTR as *Cpeb1* (Afroz et al., 2014; Wang & Tanaka Hall, 2001). These genes might thus have partially interchangeable roles, which could explain why one or the other is sometimes missing in some passerine species (Ruiz-Ruano et al., 2025). The last old gene identified on the GRC was *Pim1*, which is involved in processes such as cell cycle progression, cell proliferation and cell migration, and acts as an oncogene in many types of cancer (Nawijn et al., 2010). *Pim1* has also been identified as the only functional gene shared between the oscines and suboscines in the comparative study across passerines (Ruiz-Ruano et al., 2025). These three genes are thus the best candidates for the functional indispensability of the GRC in passerine birds.

One of the most striking features of the GRCs in *Lonchura* species is the fine-scale mosaic pattern formed by regions of different A-chromosomal origin, where younger sequences usually make larger, more continuous blocks, while older sequences are shorter and intermixed with each other. Although the composite A-chromosomal origin of GRC content has been documented previously, earlier studies interpreted this structure as the outcome of recurrent sequence additions followed by deletions (Kinsella et al., 2019; Schlebusch et al., 2023; Ruiz-Ruano et al., 2025). However, our high-quality, contiguous GRC assemblies from closely related *Lonchura* species revealed another mechanism likely contributing to GRC structural evolution. Despite sharing substantial genetic content, the *Lonchura* species exhibit extensive fine-scale within-chromosomal restructuring and a little collinearity between species (Figure 5 A). This suggests that sequences are copied from the A chromosomes in the form of relatively large intact tracts on the GRC, before being effectively diced, often duplicated, and moved around the GRC over time.

Such chromosomal instability contrasts with the conserved chromosomal structure in birds (Ellegren, 2010) and is also rarely observed in other taxa or regular chromosomes (Tan et al., 2015). However, it resembles the situation in cancer cells, where extensive and complex chromosomal rearrangements can occur due to a mutational process known as chromothripsis (Mcneil et al., 2025). Chromothripsis is often associated with the formation of micronuclei containing missegregated chromosomes. Normally, the DNA in these micronuclei is fragmented and degraded, but if the micronucleus rejoins the nucleus before this process is complete, it results in highly fragmented DNA being reintroduced to the cell’s DNA repair mechanisms to be reconnected, but in a completely different order and orientation. Since the GRC is eliminated from somatic cells and male germ cells through the formation of micronuclei, it is possible that chromothripsis may contribute to its rapid structural evolution. During GRC elimination, two GRC chromatids (unseparated during mitosis or meiosis) are likely to form one micronucleus (Dedukh et al., 2025). Their fragmentation and rejoining could therefore also explain the GRC’s propensity for duplicating sequences, similar to what has been observed with gene amplifications in tumors (Shoshani et al., 2021). Such restructuring events would normally be highly detrimental to an organism, but in the relaxed selective environment of the GRC, where rearranged and potentially damaged genes will never be expressed in the soma, this effect will be dramatically less noticeable. And in rare cases, such as observed in clitellates (Vargas-Chávez et al., 2025), massive restructuring may even be adaptive and lead to the origin of new lineages.

Together, our analyses of the currently highest-quality GRC assemblies reveal that the GRC is an exceptionally dynamic and rapidly evolving component of an otherwise relatively stable passerine genome, with substantial structural changes occurring even on very recent evolutionary time scales. We demonstrate for the first time that this fast structural evolution is driven not only by extensive sequence turnover, but also by chromothripsis-like processes likely associated with GRC elimination. In addition, we revealed that rapid and enormous changes in GRC sizes among species are caused by massive amplification of different satellite sequences likely enabled by relaxed selection pressures on the GRC. Thus, the GRC’s accelerated evolution is driven not only by altered selection pressures caused by restriction of the chromosome to the germline, but also by specific mutational processes associated with chromosome elimination mechanisms. The GRC may therefore constitute a unique genomic environment that allows duplicated genomic material to accumulate, be reshuffled, and potentially acquire novel functions, thereby facilitating evolutionary innovation or even contributing to speciation. Remarkably, despite this highly dynamic background, a small set of ancient genes has persisted on the GRC for tens of million years. Notably, two of these genes, *Cpeb1* and *Elavl4*, are involved in regulating gene expression on the translational level and may play key roles in establishing differential gene expression programmes between germline and soma. Their persistence on the GRC may explain why the GRC has not been lost but has instead become an indispensable component of the passerine genome.

The selective advantages conferred by such genome compartmentalization may also account for the repeated, independent evolution of GRCs in other taxa, including lampreys and hagfish (Smith et al., 2021), as well as dipteran flies (Hodson & Ross, 2021). In all these lineages, GRCs have been maintained for tens of million of years, and despite having some taxon-specific features, they share striking similarities in elimination mechanisms and patterns of GRC evolution (Hodson et al., 2025; Kerrebrock et al., 2025; Marlétaz et al., 2024; Smith et al., 2018; Timoshevskiy et al., 2016). GRCs may therefore represent a relatively widespread yet underappreciated driver of genome evolution in animals.

## Methods

### Sample collection and DNA extraction

One male individual from each of the four *Lonchura* species (*L. malacca, L. punctulata, L. castaneothorax* and *L. domestica*) was sampled for whole genome sequencing. The birds were euthanized by cervical dislocation and had a left testis and a kidney dissected, frozen in liquid nitrogen, and stored at −80 °C for DNA extraction. High molecular weight DNA was isolated following a phenol-chloroform procedure as described in Pajer et al., (2006), except that the tissues were incubated in the lysis buffer for several days (until mostly dissolved) and 2 volumes of 80% ethanol were used for the DNA precipitation. In addition, the right testis was dissected from *L. punctulata* and *L. malacca* individuals, frozen in liquid nitrogen and shipped on dry ice to the Institute of Applied Biotechnologies (Olomouc, Czech Republic) for Dovetail Omni-C sequencing.

### Sequencing

The high molecular weight DNA extracted from testes and kidneys was sequenced using PacBio High Fidelity (HiFi) sequencing on the PacBio Revio system at the National Genomics Infrastructure (NGI, Uppsala, Sweden). Due to the fact that the GRC is only found in germline cells within the testis, and it is hemizygous in those cells, the expected depth is a lot lower than the expected depth across A chromosomes. Therefore, in order to ensure sufficient depth across the GRC, each testis was sequenced on a full Revio SMRT Cell, while two kidney samples were pooled together before being sequenced on a single SMRT Cell. This resulted in an average HiFi depth of 74x for the testis samples and 31x depth for the kidney samples, assuming a genome size of 1.1 Gbp (see Supplementary Table 2). The N50 of these samples ranged between 12-18 kbp while the N90 was 6-14 kbp. The *L. malacca* and *L. punctulata* Omni-C libraries (from the individual’s other testis) were sequenced once with a read length of 150 bp (using the Illumina NovaSeq X instrument) and once with a read length of 300 bp (using the Illumina NextSeq 2000 instrument).

### Assembly and GRC identification

The kidney genomes were assembled for each species with hifiasm v0.20.0-r639 (Cheng et al., 2024) on the default setting. The testis genomes were also assembled with hifiasm, but with merging switched off (-m 0) and Omni-C data included in the case of *L. punctulata* and *L. malacca*. The testis and kidney PacBio reads were then aligned to the testis assemblies using minimap2 v 2.26-r1175 (Li, 2021). The read depth across each contig was calculated for both kidney and testis datasets and normalised according to the average depth across the testis genome.

An initial lenient selection of possible GRC contigs was selected where the normalised testis read depth was 1.5x greater than the normalised kidney read depth. The filter of only requiring 1.5x higher normalised depth across a contig was deliberately lenient in order to not lose any GRC contigs. The selected contigs were then inspected and further refined manually to exclude potential A-chromosomal contigs as described below.

In the case of the micro GRCs (*L. punctulata* and *L. malacca*), Omni-C data and the implied spatial proximity was an important consideration when deciding which contigs to include and exclude in the final list of GRC contigs. However, the mismapping of A-chromosomal reads onto recently duplicated GRC sequences risked adding noise to the analysis. In order to minimise the effects of this mismapping, the frequency of 35-bp kmers was calculated across the testis assemblies using jellyfish v2.3.1 (Marçais & Kingsford, 2011). A selection of kmers with a frequency two or more was inverted to select portions of the testis genome with a unique sequence (a frequency of one). Omni-C reads were aligned to the *L. punctulata* and *L. malacca* genomes using bwa mem v0.7.17-r1188 and the “-Sw” option (Li, 2013). Read pairs were then discarded which didn’t have both reads touch a unique region of the testis assembly (see Supplementary Figure 12 and 13).

The initial set of selected GRC contigs was then characterized by other criteria: first, each putative GRC sequence had the testis and kidney normalised PacBio read depth calculated in windows of 10 kbp, with each window starting 2 kbp along the sequence, and no window being smaller than 5 kbp. Second, these windows had the frequency within the putative GRC of 35-bp kmers quantified. This gives a measure of the extent any contigs’ sequence was found in the rest of the GRC. Third, the windows had the proportion of the 35-bp kmers which were not found in the kidney dataset quantified. Finally, each putative GRC contig was aligned to the *T. guttata* somatic genome using minimap2 (accession number GCF_048771995.1), discarding alignments less than 1 kbp, in order to identify the A-chromosomal origin.

Once these factors were viewed together with the Omni-C data’s connections, it was possible to distinguish real GRC sequences showing consistent characteristics (see Results) from sequences that were prone to be erroneously added to the GRC due to the lenient testis/kidney read depth filter. We excluded contigs if they had unusual testis/kidney read depth, kmers that were not found on the other GRC contigs, too few kmers that were only found in the testis dataset, if they had a single A-chromosomal origin, as well as, in the case of L. malacca and L. punctulata, a lack of Omni-C connections.

### Estimation of the age of GRC sequences

After aligning each GRC to the *T. guttata* genome using minimap2, the GRC sequence from each alignment greater than 1 kb was further split up into windows, a maximum of 2.5 kb and a minimum of 1 kb, starting every 500 bp. These smaller windows were then aligned to the 4 *Lonchura* somatic genome assemblies made from the kidney data as well as somatic genome assemblies from *T. guttata*, *Luscinia megarhynchos* (accession GCA_034336665.1), *Chiroxiphia lanceolata* (GCF_009829145.1) and *Falco peregrinus* (GCF_023634155.1).

Each window that had 80% of its sequence aligned to every genome was selected along with the genomic sequences they aligned to and aligned together using muscle v5.2 (Edgar, 2004). The multiple sequence alignments were then used to create a genealogy using FastTree v2.2.0 and the ‘gtr + cat’ model (Price et al., 2010). These trees were then rooted using *F. peregrinus*. Each resultant tree was classified according to where the GRC sequence diverged from the A-chromosomal sequences. If the genealogy within the Lonchura genus did not correspond to the expected phylogeny (a possible result of ancestral polymorphism or hybridization among the species) the GRC sequence was classified as diverging somewhere within the clade. In cases where the genealogy did not correspond to phylogeny in other than *Lonchura* species, the genealogy was considered unclassifiable and discarded.

Individual minimap2 alignments (that correspond to sections of GRC of the same A-chromosomal origin) were therefore composed of one or more windows which had been classified as to when that GRC sequence started to diverge from the A-chromosomal paralog. Most alignments had a clear majority consensus from among the windows as to when the GRC sequence started to diverge. In these cases, the whole alignment was then given this classification. Cases where there was no majority consensus were relatively rare and the classification was done manually by merging the time points to express the uncertainty.

### Repeat content analysis

A repeat library was built for each of the four species. First, a *de novo* repeat library was constructed by RepeatModeler (v.2.0.7) (Flynn et al., 2020) with default settings using a) A-chromosomal and GRC sequence together, b) only GRC sequence. Second, a long terminal repeat retroelements (LTR) library was built by combining the outputs of LTRharvest (part of GenomeTools v.1.6.5, parameters -minlenltr 100 -maxlenltr 7000 -mintsd 4 -maxtsd 6 -motif TGCA -motifmis 1 -similar 85 -vic 10 -seed 20 -seqids yes) (Ellinghaus et al., 2008) and LTR_FINDER (v.1.3, parameters -threads 10 -harvest_out -size 1000000 -time 300) (Xu & Wang, 2007) using LTR_retriever (v.2.9.0) (Ou & Jiang, 2018). These programs used the concatenation of A-chromosomal and GRC sequence as their input. And finally, python script famdb.py (v.2.0.5, part of RepeatMasker distribution) (Smit et al., 2013) was used to create a taxon-specific subset of the Dfam database (v.3.9) (Storer et al., 2021) - more specifically, repeat families ancestral and descendent to taxa *Lonchura* (ID 40156) and *T. guttata* (ID 59729) were extracted. Species-specific repeat libraries were then built by concatenation of these four libraries for each species.

Tandem repeat finder (TRF) (v.4.09) (Benson, 1999) with recommended settings was used to explore the satellite profiles within the four GRCs. For *L. domestica*, a dimer of consensus sequence of the most abundant satellite from the TRF was blasted (BLASTN, v.2.17.0+, (Morgulis et al., 2008; Zhang et al., 2000) against the NCBIcore_nt database resulting in a hit (E-value 1e-39, 83% sequence identity) with Tgut191A sequence (accession number ON037476.1) (Supplementary Figure 6), published centromere-associated repeat from *T. guttata* (Takki et al., 2022). For *L. castaneothorax*, a consensus sequence of the 17-bp satellite was derived manually from the consensus sequences of the most common 17-bp satellites reported by TRF (ACGAGCTGGGTAAGGAC) and called Lcas17GRC.

These satellite sequences (Tgut191A and Lcas17GRC) were masked in the A-chromosomal and GRC assemblies using RepeatMasker (v.4.2.2) (Smit et al., 2013). Subsequently, in a second RepeatMasker round, species-specific repeat libraries were used to analyze the remaining repeat content.

### Gene annotation and functionality analysis

Genes were annotated on the four GRCs by aligning the proteins from *T. guttata* to each of them using Exonerate v2.4.0 (--model protein2genome --percent 50 --showcigar yes --showquerygff yes --showtargetgff yes --minintron 10) (Slater & Birney, 2005). When multiple proteins’ exons aligned to the same place on a GRC, only the protein with the best alignment score was annotated. We then identified the location of stop codons, frameshift mutations, deletions, and insertions, as well as the location and size of introns, using in-house Python scripts. Afterwards, a gene was considered functional if more than 90% of its expected length was found without any premature stop codons.

The location and size of introns were also registered for *T. guttata*, which was used as reference in the comparisons performed to the GRCs. To infer gene duplication events, we used the gene coding sequence, and created multiple sequence alignments with MAFFT v7.520 (Katoh & Standley, 2013). After this, we refined the alignments by removing uninformative sites using Clipkit v.2.7.0 (Steenwyk et al., 2020), nucleotide positions represented in less than 70% of sequences, and sequences with information in fewer than 50% of aligned sites. The resulting alignments were then used to construct maximum likelihood phylogenetic trees with IQ-TREE v.3.0.1 (Kalyaanamoorthy et al., 2017; Hoang et al., 2017; Wong et al., 2025), using 10,000 ultrafast bootstrap replicates. To finalize, we visualized and edited the trees with TreeViewer v.2.2.0 (Bianchini & Sánchez-Baracaldo, 2024). The alignment and genealogy were used to estimate gene duplication events with MEGA v.12.1.1 (Kumar et al., 2024).

### GO term enrichment

Gene Ontology (GO) enrichment analysis was performed using ClusterProfiler (enrich) (Yu et al., 2012) with the 15,020 annotated *T. guttata* genes as background. For KEGG pathway enrichment analysis we used ShinyGO v.0.85 (Ge et al., 2020; Kanehisa et al., 2021). Significant results were filtered using a false discovery rate of 0.05.

### GRC-linked gene genealogies

The genealogies of individual GRC genes were done by extracting A-chromosomal paralogs of GRC-linked genes for 13 avian species using Exonerate as described above. For each gene, the sequence with the best alignment score was selected. In order to avoid tree reconstruction biases, in GRC-linked multicopy genes, a single sequence was selected for each species. Preference was given to functional sequences when available. The alignments and genealogies were created using maximum likelihood phylogenetic approach as described above in Gene annotation and functionality analysis; the only change was filtering out codons that were unreliable or represented in less than 70% of sequences, rather than using the same criteria for nucleotide positions.

## Acknowledgements

We thank the GRC Brainstorming group and members of the Radka Reifova’s and Alexander Suh’s labs for helpful discussions, especially Francisco Ruiz-Ruano. The work was funded by the Czech Science Foundation (grant 23-07287S to R.R., D.D., T.A. and grant 25-17195S to R.R. and J.P.), the Grant Agency of Charles University (GAUK 314222 to Z.H.) and SVV 260818/2025, a Consolidator Grant of the European Research Council (101002158 GermlineChrom to A.S.) and a Project Grant of the Swedish Research Council Vetenskapsrådet (2020-04436 to A.S.). Computational analysis was done with the Institute of Molecular Genetics (Czech Academy of Sciences, Prague, Czech Republic) computers with support from ELIXIR CZ Research Infrastructure (ID LM2023055, MEYS CR).

## Data availability

Raw sequence data and GRC assemblies have been uploaded to NCBI (bioproject PRJNA1400222). GRC assemblies have also been uploaded onto figshare (10.6084/m9.figshare.31343230).

## Notes

### Competing Interest Statement

The authors have declared no competing interest.

