## Supplementary Material for "Stable but turbulent: the two faces of the germline-restricted chromosome of passerine birds"

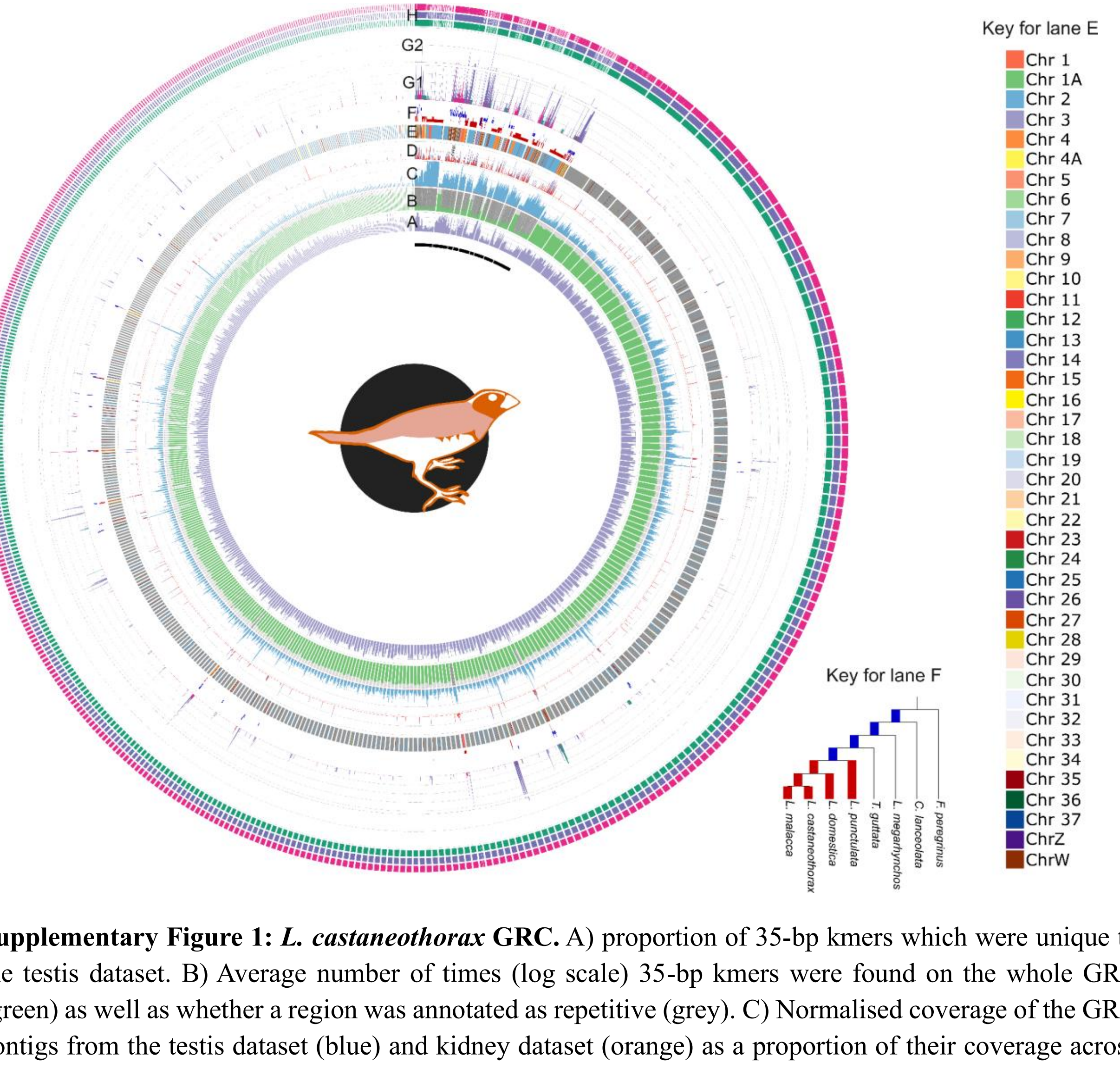


**Supplementary Figure 1:** ***L. castaneothorax*** **GRC.** A) proportion of 35-bp kmers which were unique to the testis dataset. B) Average number of times (log scale) 35-bp kmers were found on the whole GRC (green) as well as whether a region was annotated as repetitive (grey). C) Normalised coverage of the GRC contigs from the testis dataset (blue) and kidney dataset (orange) as a proportion of their coverage across the whole genome. D) Gene density for all identified genes (purple) as well as only functional genes (red). E) A chromosome that the GRC sequence aligned to. F) Estimated age of the GRC region. The young sequence (red) originated after the diversification of the *Lonchura* species (~4 mya) while the older sequence (blue) predates the diversification. If there was ambiguity about whether a sequence diverged from the A chromosomes before or after the Lonchura diversification, it is shown in purple. The further away from the center of the Circos plot, the older the region (see Key for lane F). G) Alignment of the other *Lonchura* species' GRCs to *L. malacca*. This is shown on a linear scale (G1) for less than or equal to 8 alignments, after which it is shown on a log scale (G2). H) Indicates whether the *L. castaneothorax* GRC aligned to *L. malacca* (green), *L. domestica* (purple), and *L. punctulata* (pink). Lanes A, B, C, and D used a sliding window of 10 kbp and a step size of 2 kbp (minimum size 5 kbp). Lane F and G used a sliding window of 2.5 kbp and a step size of 500 bp (minimum size 1 kbp). The first 9 scaffolds are highlighted with a black line underneath them. These scaffolds were used in Figure 5 and can be seen in more detail in Supplementary Figure 3.

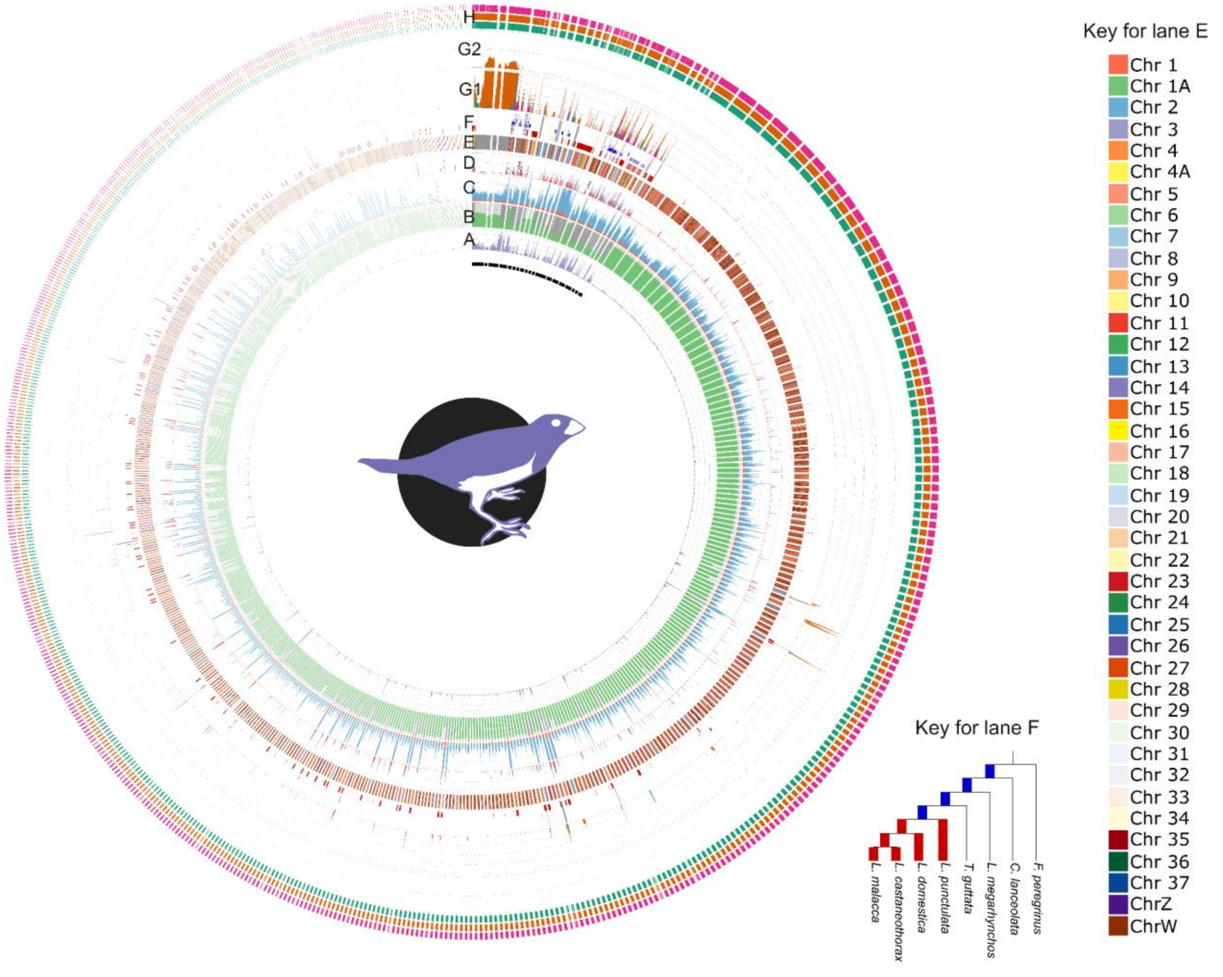


**Supplementary Figure 2:** ***L. domestica*** **GRC.** A) proportion of 35-bp kmers which were unique to the testis dataset. B) Average number of times (log scale) 35-bp kmers were found on the whole GRC (green) as well as whether a region was annotated as repetitive (grey). C) Normalised coverage of the GRC contigs from the testis dataset (blue) and kidney dataset (orange) as a proportion of their coverage across the whole genome. D) Gene density for all identified genes (purple) as well as only functional genes (red). E) A chromosome that the GRC sequence aligned to. F) Estimated age of the GRC region. The young sequence (red) originated after the diversification of the *Lonchura* species (~4 mya) while the older sequence (blue) predates the diversification. If there was ambiguity about whether a sequence diverged from the A chromosomes before or after the Lonchura diversification, it is shown in purple. The further away from the center of the Circos plot, the older the region (see Key for lane F). G) Alignment of the other *Lonchura* species' GRCs to *L. malacca*. This is shown on a linear scale (G1) for less than or equal to 8 alignments, after which it is shown on a log scale (G2). H) Indicates whether the *L. domestica* GRC aligned to *L. malacca* (green), *L. castaneothorax* (orange), and *L. punctulata* (pink). Lanes A, B, C, and D used a sliding window of 10 kbp and a step size of 2 kbp (minimum size 5 kbp). Lane F and G used a sliding window of 2.5 kbp and a step size of 500 bp (minimum size 1 kbp). The first 19 scaffolds are highlighted with a black line underneath them. These scaffolds were used in Figure 5 and can be seen in more detail in Supplementary Figure 4.

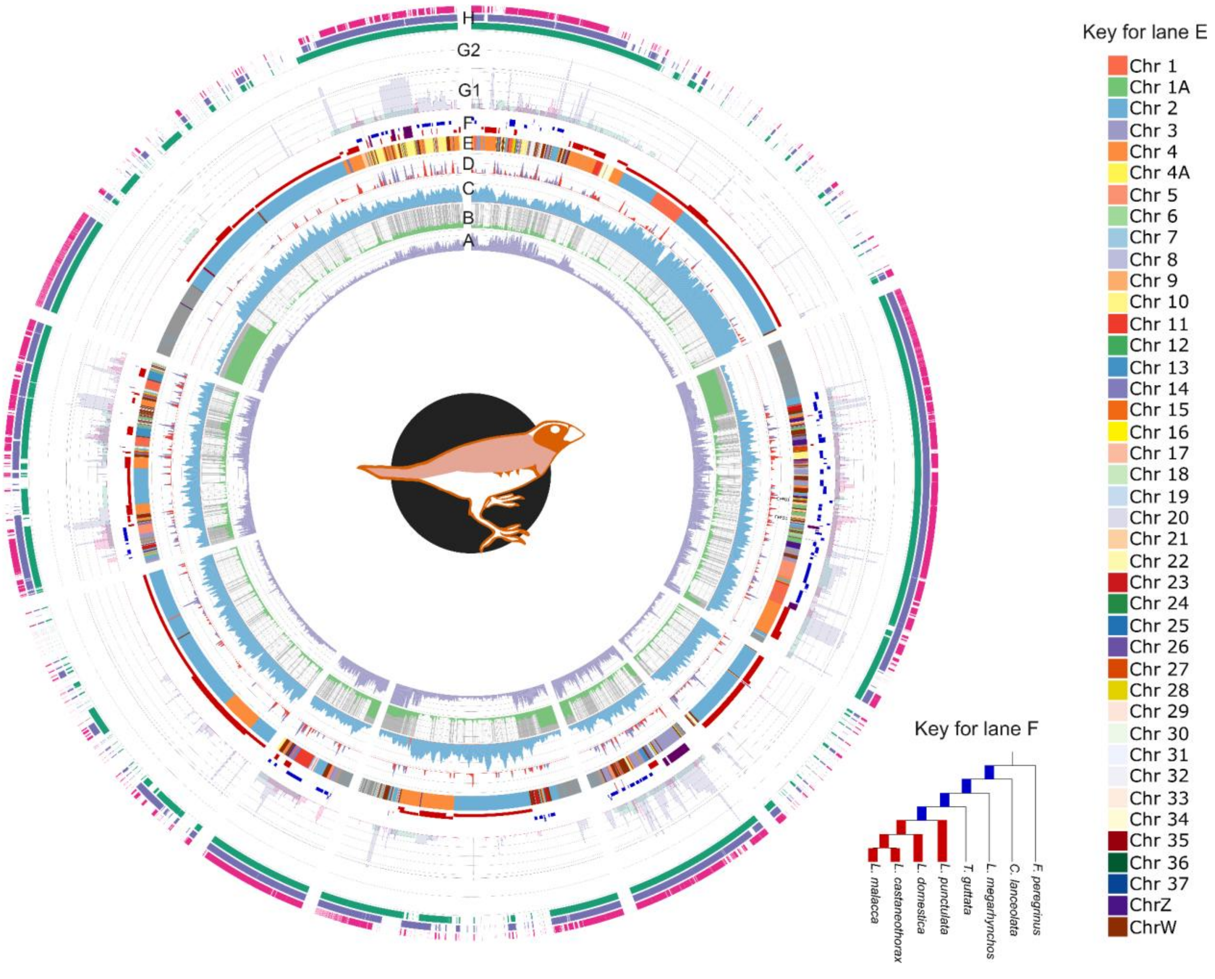


**Supplementary Figure 3: Subset of the *L. castaneothorax* GRC.** A) proportion of 35-bp kmers which were unique to the testis dataset. B) Average number of times (log scale) 35-bp kmers were found on the whole GRC (green) as well as whether a region was annotated as repetitive (grey). C) Normalised coverage of the GRC contigs from the testis dataset (blue) and kidney dataset (orange) as a proportion of their coverage across the whole genome. D) Gene density for all identified genes (purple) as well as only functional genes (red). E) A chromosome that the GRC sequence aligned to. F) Estimated age of the GRC region. The young sequence (red) originated after the diversification of the *Lonchura* species (~4 mya) while the older sequence (blue) predates the diversification. If there was ambiguity about whether a sequence diverged from the A chromosomes before or after the Lonchura diversification, it is shown in purple. The further away from the center of the Circos plot, the older the region (see Key for lane F). G) Alignment of the other *Lonchura* species' GRCs to *L. malacca*. This is shown on a linear scale (G1) for less than or equal to 8 alignments, after which it is shown on a log scale (G2). H) Indicates whether the *L. castaneothorax* GRC aligned to *L. malacca* (green), *L. domestica* (purple), and *L. punctulata* (pink). Lanes A, B, C, and D used a sliding window of 10 kbp and a step size of 2 kbp (minimum size 5 kbp). Lane F and G used a sliding window of 2.5 kbp and a step size of 500 bp (minimum size 1 kbp). The first 9 scaffolds are highlighted with a black line underneath them. These scaffolds were used in Figure 5 and can be seen in more detail in Supplementary Figure 3.

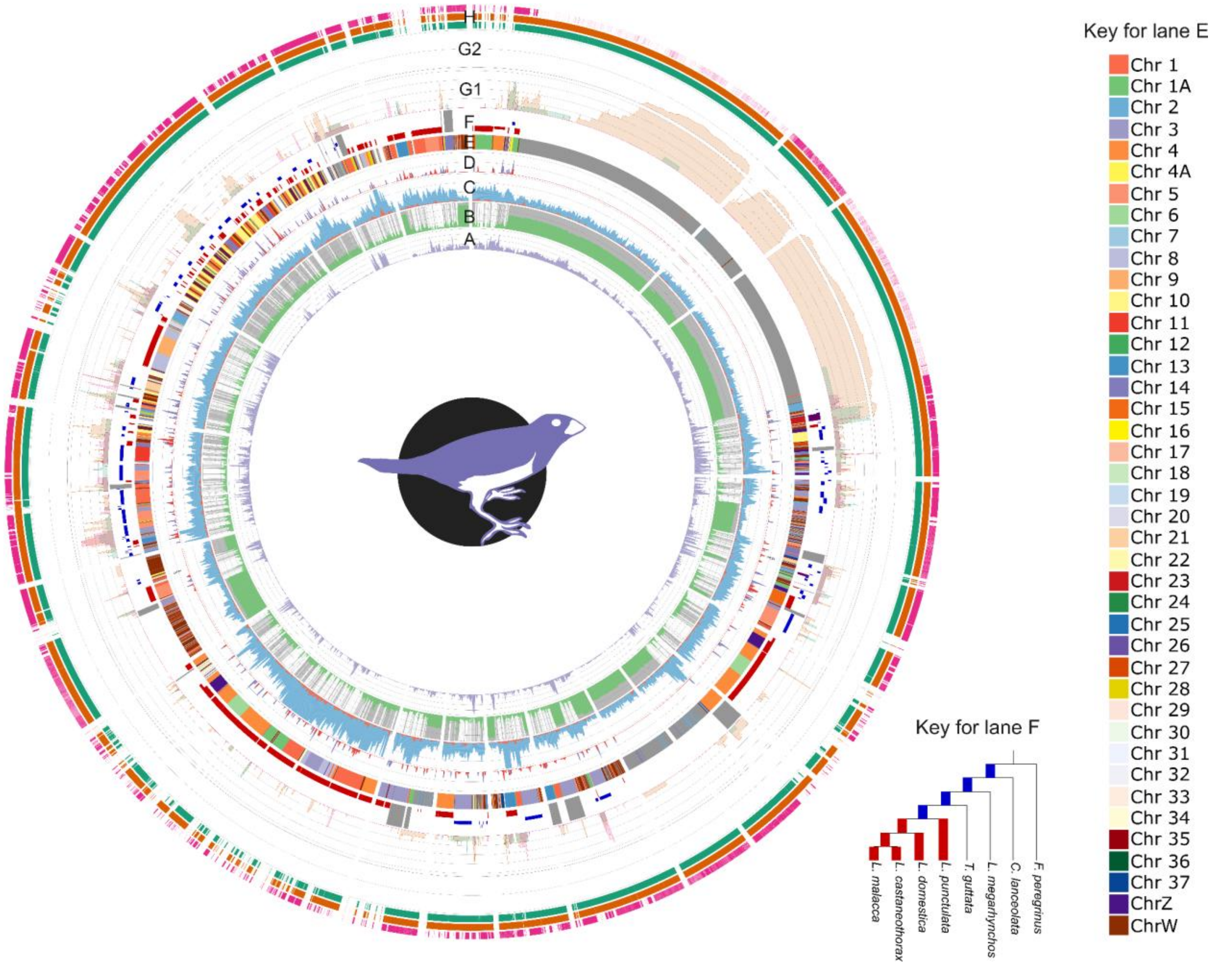


**Supplementary Figure 4: Subset of the *L. domestica* GRC.** A) proportion of 35-bp kmers which were unique to the testis dataset. B) Average number of times (log scale) 35-bp kmers were found on the whole GRC (green) as well as whether a region was annotated as repetitive (grey). C) Normalised coverage of the GRC contigs from the testis dataset (blue) and kidney dataset (orange) as a proportion of their coverage across the whole genome. D) Gene density for all identified genes (purple) as well as only functional genes (red). E) A chromosome that the GRC sequence aligned to. F) Estimated age of the GRC region. The young sequence (red) originated after the diversification of the *Lonchura* species (~4 mya) while the older sequence (blue) predates the diversification. If there was ambiguity about whether a sequence diverged from the A chromosomes before or after the Lonchura diversification, it is shown in purple. The further away from the center of the Circos plot, the older the region (see Key for lane F). G) Alignment of the other *Lonchura* species' GRCs to *L. malacca*. This is shown on a linear scale (G1) for less than or equal to 8 alignments, after which it is shown on a log scale (G2). H) Indicates whether the *L. domestica* GRC aligned to *L. malacca* (green), *L. castaneothorax* (orange), and *L. punctulata* (pink). Lanes A, B, C, and D used a sliding window of 10 kbp and a step size of 2 kbp (minimum size 5 kbp). Lane F and G used a sliding window of 2.5 kbp and a step size of 500 bp (minimum size 1 kbp). The first 19 scaffolds are highlighted with a black line underneath them. These scaffolds were used in Figure 5 and can be seen in more detail in Supplementary Figure 4.

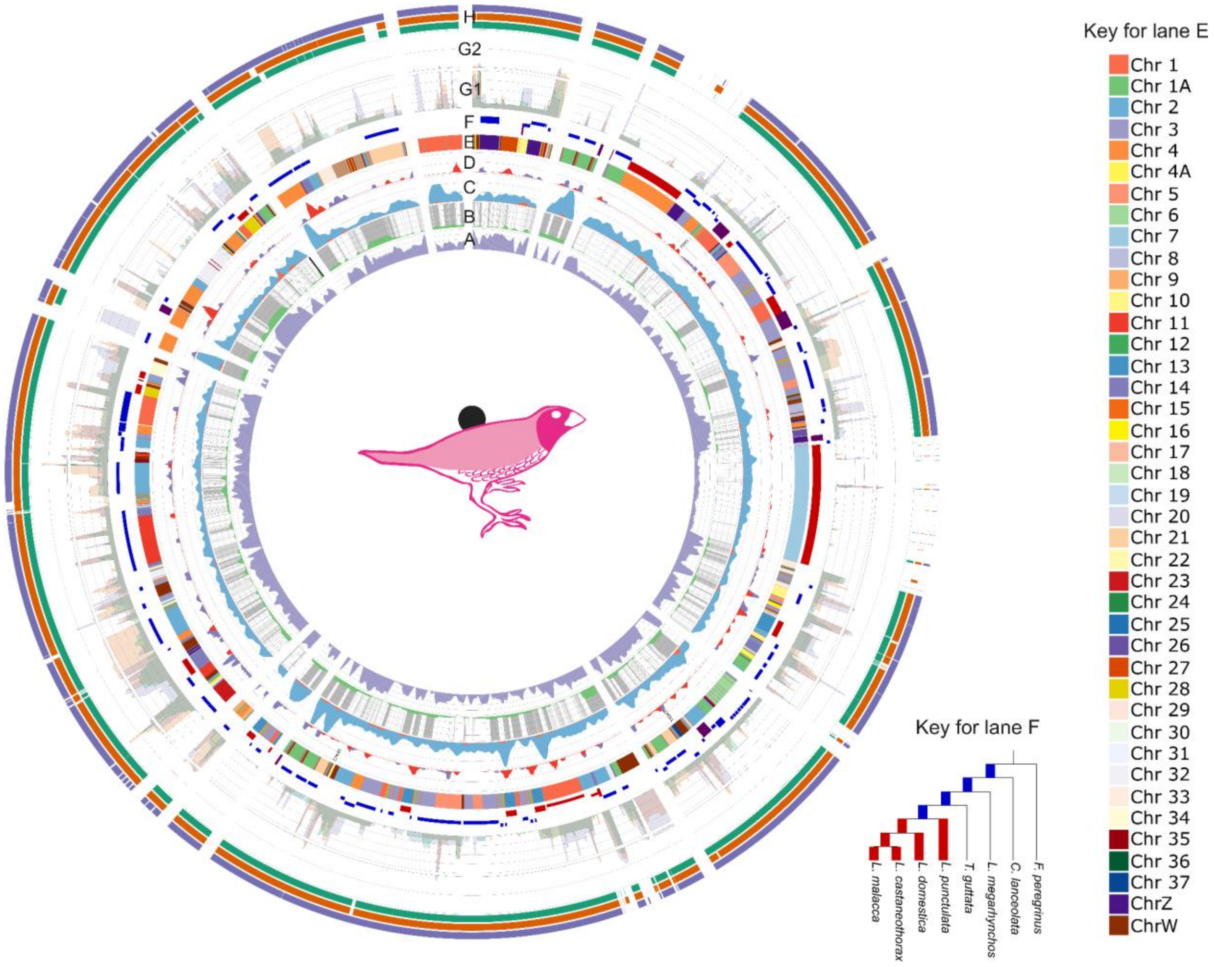


**Supplementary Figure 5:** ***L. punctulata*** **GRC.** A) proportion of 35-bp kmers which were unique to the testis dataset. B) Average number of times (log scale) 35-bp kmers were found on the whole GRC (green) as well as whether a region was annotated as repetitive (grey). C) Normalised coverage of the GRC contigs from the testis dataset (blue) and kidney dataset (orange) as a proportion of their coverage across the whole genome. D) Gene density for all identified genes (purple) as well as only functional genes (red). E) A chromosome that the GRC sequence aligned to. F) Estimated age of the GRC region. The young sequence (red) originated after the diversification of the *Lonchura* species (~4 mya) while the older sequence (blue) predates the diversification. If there was ambiguity about whether a sequence diverged from the A chromosomes before or after the Lonchura diversification, it is shown in purple. The further away from the center of the Circos plot, the older the region (see Key for lane F). G) Alignment of the other *Lonchura* species' GRCs to *L. malacca*. This is shown on a linear scale (G1) for less than or equal to 8 alignments, after which it is shown on a log scale (G2). H) Indicates whether the *L. punctulata* GRC aligned to *L. malacca* (green), *L. castaneothorax* (orange), and *L. domestica* (purple). Lanes A, B, C, and D used a sliding window of 10 kbp and a step size of 2 kbp (minimum size 5 kbp). Lane F and G used a sliding window of 2.5 kbp and a step size of 500 bp (minimum size 1 kbp).

```
Query: LDom_TRF_consensus_sequence Query ID: lcl|Query_4968055 Length: 191


>Taeniopygia guttata satellite Tgut191A sequence
Sequence ID: ON037476.1 Length: 191
Range 1: 1 to 191

Score:176 bits(95), Expect:1e-39,
Identities:159/191(83%),  Gaps:0/191(0%), Strand: Plus/Plus

Query  1    AAACTGACATCAGGAACCTGGACTGGTGAGTTTTCTTTCCCAGGGAAAATGGCCAAAGGA  60
            |||||||||||||||||||||||||||||||||||||||||||||||||  ||| | |  
Sbjct  1    AAACTGACATCAGGAACCTGGACTGGTGAGTTTTCTTTCCCAGGGAAAACAGCCGATGTT  60

Query  61   TGTGCCCAGGGGCCCCTGGCACTGGGCTGGGACCTGACTGCAAAAAATCCTTATATCGAA  120
            ||||| ||||| ||  |||| ||| || || | |||||||||||||||||  | ||||||
Sbjct  61   TGTGCTCAGGGCCCTGTGGCTCTGAGCCGGCAGCTGACTGCAAAAAATCCGAACATCGAA  120

Query  121  GGATGGACCCCAGACCGAAGTTGTCCCCTCCTCAAACTTGTGCTGCCCTGATAGACCCAA  180
            ||||    ||||||| | || || |||| ||||| |||||||||||||||| || |||| 
Sbjct  121  GGATAAGTCCCAGACAGGAGCTGCCCCCACCTCATACTTGTGCTGCCCTGACAGTCCCAG  180

Query  181  GGGAGCCCTAC  191
             ||||||||||
Sbjct  181  AGGAGCCCTAC  191
```

**Supplementary Figure 6: Alignment of the 191 bp *Lonchura domestica* GRC repeat.** The repeat (cumulatively forming 74% of the assembled GRC sequence) is aligned to Tgut191A, a centromere-associated tandem repeat from *Taeniopygia guttata*.

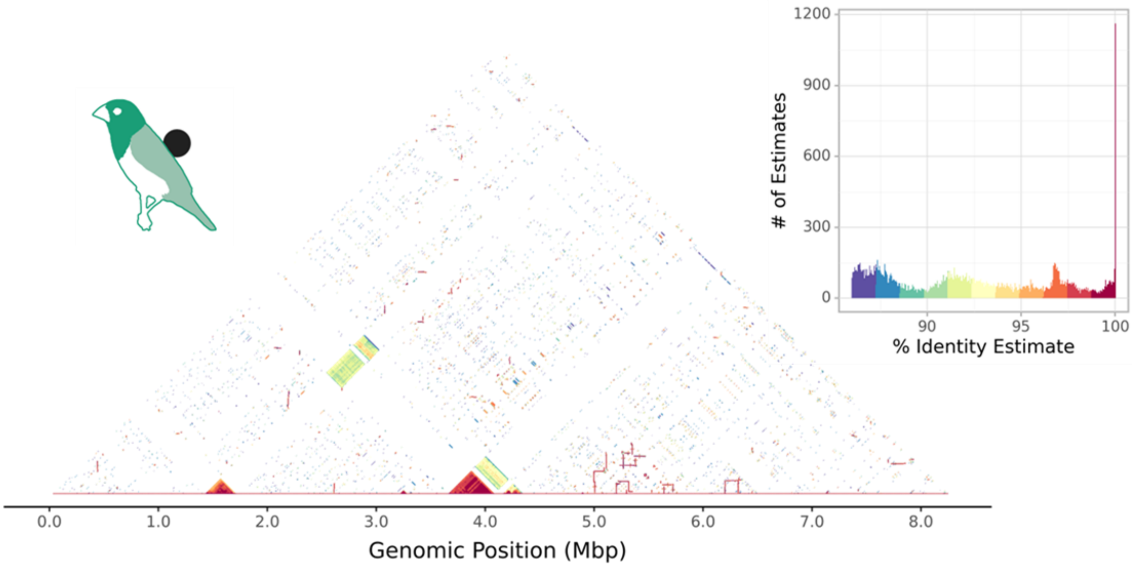


**Supplementary Figure 7: Alignment of the *L. malacca* GRC to itself to identify repeat sequences.** Two main repeat structures are clear, one between at about 1.5 Mbp along the chromosome and the other at 4

Mbp along. Each repeat structure has a high percentage identity match to itself (see the key in the top right) and low homology with the other repeat.

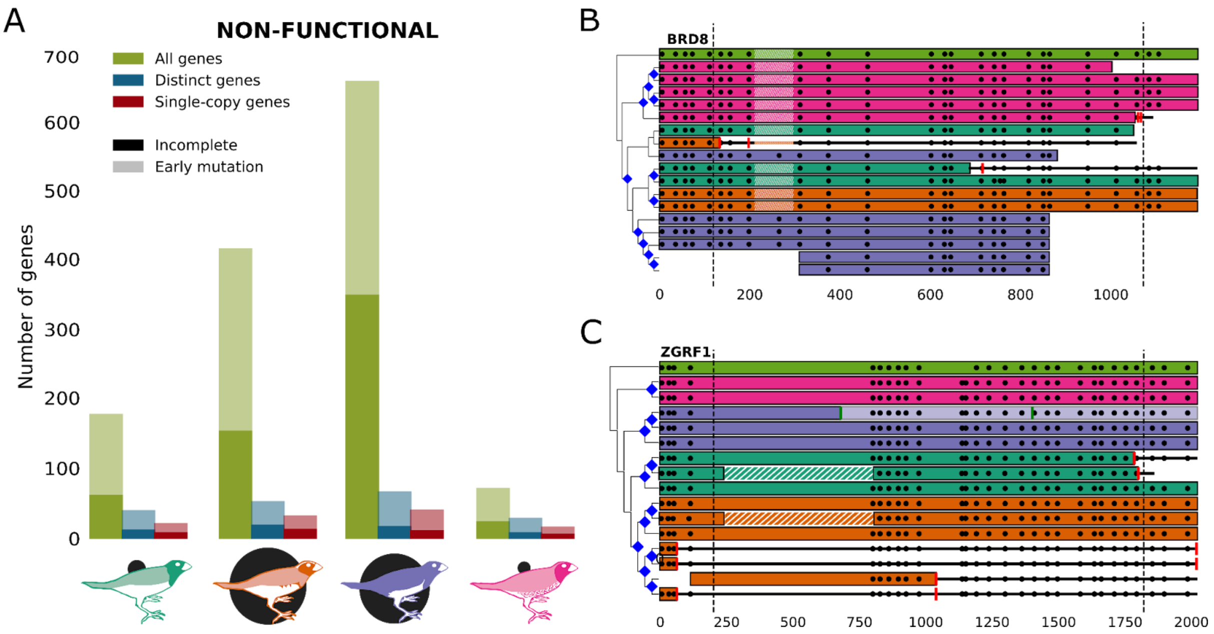


**Supplementary Figure 8: Reasons for non-functional classification.** A) The number of genes per species classified as non-functional and whether that was due to incomplete coding sequence or the presence of premature nonsense or frame shift mutations. B-C) Representation of selected gene structures showing pseudogenization events. Genes are shown as horizontal bars, coloured according to their species, with exon boundaries represented by black dots. Frameshift mutations are represented by a vertical green line across a gene, while red lines represent a stop codon. Sequences are ordered from top to bottom according to the genealogy on their left, with the *T. guttata* A-chromosomal paralog as the outgroup. Genes that match a shorter *T.guttata* protein isoform have striped bars that illustrate the region not present in that alternative isoform. Blue lozenges in nodes of the genealogy represent predicted duplication events.

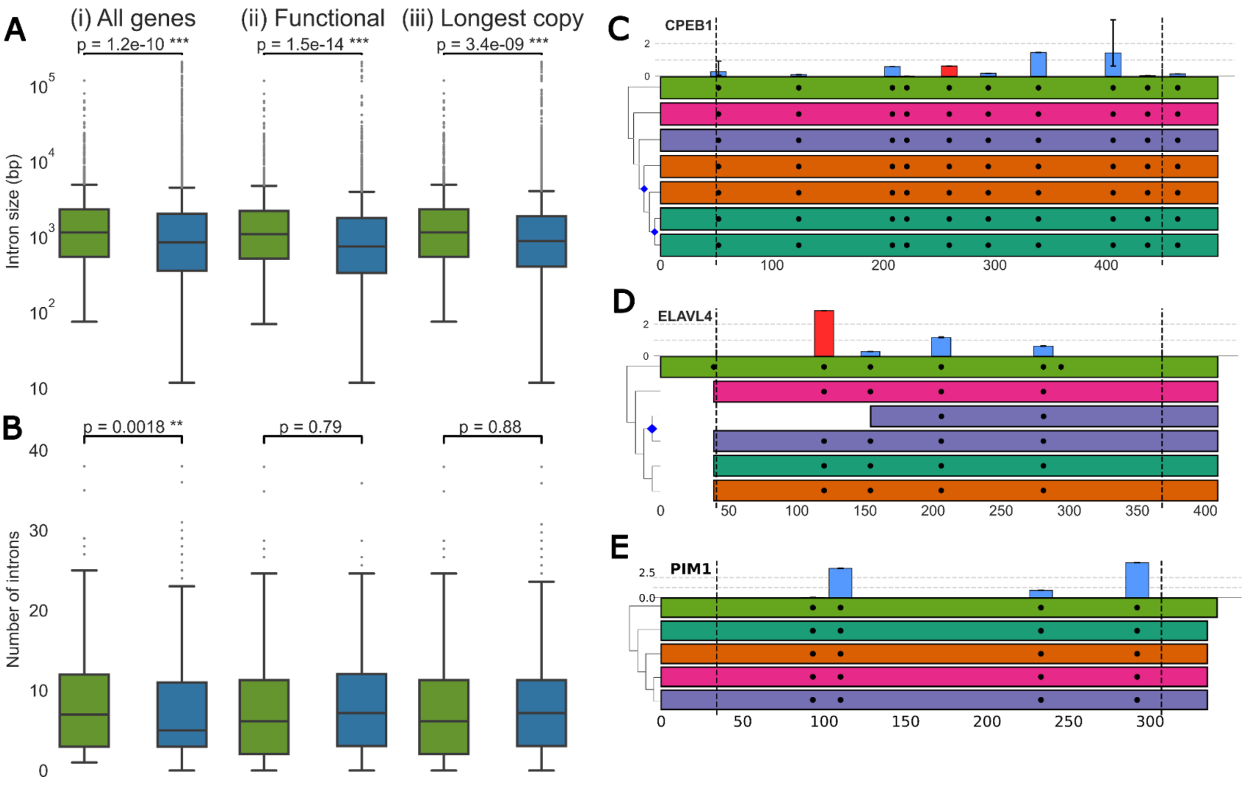


**Supplementary Figure 9: Structural evolution of GRC genes.** A) Distribution and statistical comparison of intron lengths of GRC genes (blue) and their *T. guttata* A-chromosomal paralogs (green). The comparison is shown for: all genes including non-functional ones (i), functional genes only (ii), and the longest gene copy in each species (iii). This comparison only included equivalent intron comparisons to avoid any bias from the shortened GRC genes. B) Distribution and statistical comparison of intron number of GRC genes (blue) and their *T. guttata* A-chromosomal paralogs (green). The comparison is shown for: all genes including non-functional ones (i), functional genes only (ii), and the longest gene copy in each species (iii). This comparison only included equivalent intron comparisons to avoid any bias from the shortened GRC genes. C-E) Representation of selected gene structures showing pseudogenization events and intron size changes. Genes are shown as horizontal bars, coloured according to their species, with exon boundaries represented by black dots. Frameshift mutations are represented by a vertical green line across a gene, while red lines represent a stop codon. Sequences are ordered from top to bottom according to the genealogy on their left, with the T. guttata A-chromosomal paralog as the outgroup. Blue lozenges in nodes of the genealogy represent predicted duplication events. On top of the A-chromosomal gene and for each intron position, is shown the average fold change (log2 scale) between the intron size in GRCs and the A-chromosomal paralog. Bars for shortened GRC introns (negative fold change) are red, while bars for enlarged GRC introns (positive fold change) are blue. Within-GRC variation is illustrated by the error bars showing the maximum and minimum values.

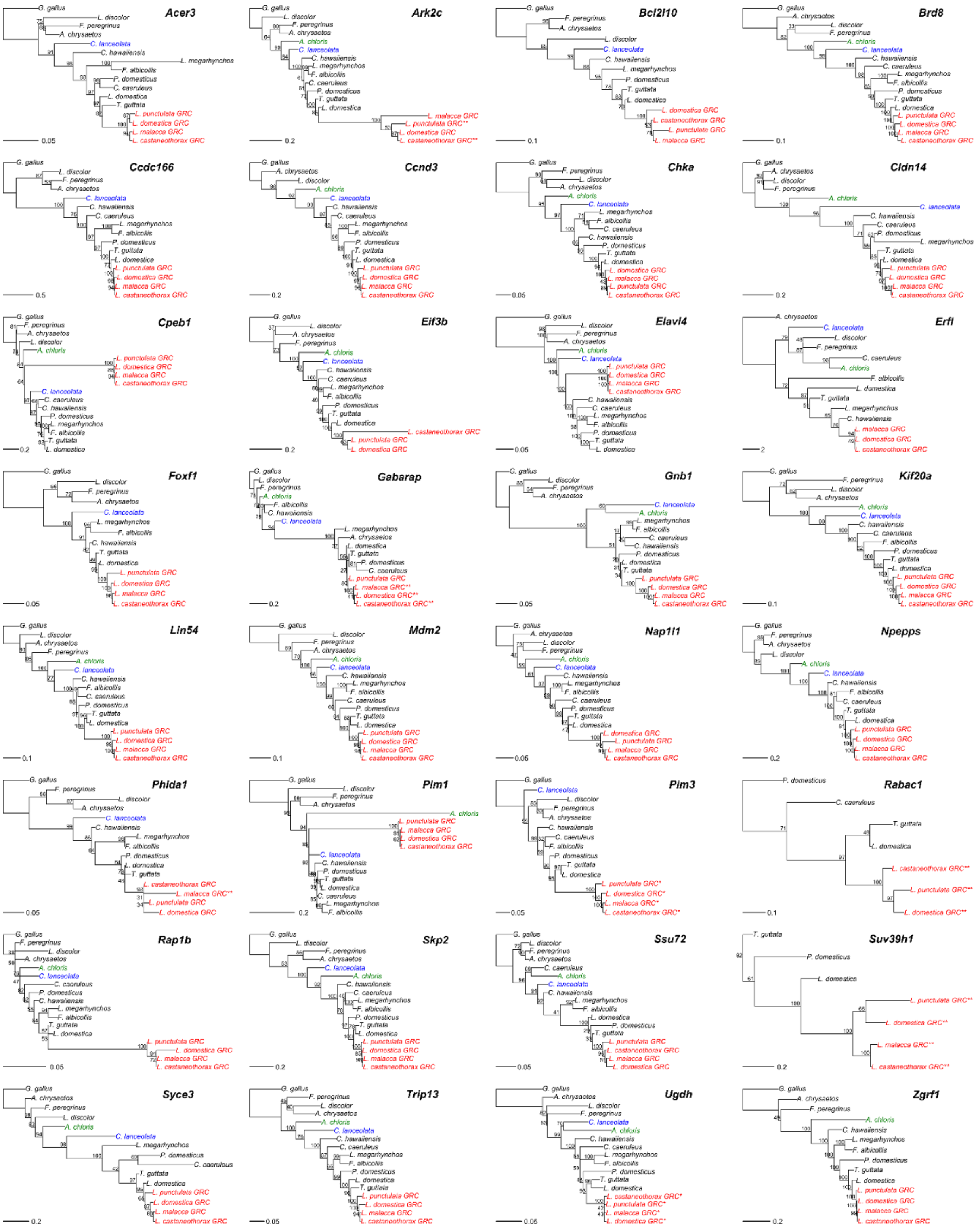


**Supplementary Figure 10: Genealogies of shared genes.** Maximum likelihood phylogenetic trees for the only thirty-two genes shared by the four GRCs and with a functional copy in at least 3 of them. GRC genes are highlighted in red, the suboscine sequence in blue, and the Acanthisitta chloris sequence in green.

Bootstrap support values are indicated at the nodes. * sequence with a nonsense mutation in the last 90% of its expected length; ** incomplete sequence (90% or less of the expected size).

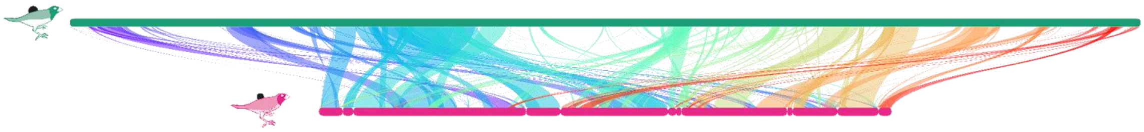

**Supplementary Figure 11: Collinearity plot of *L. malacca*'s (green) and *L. punctulata*'s (pink) GRC.** Contigs of *L. punctulata* were ordered and oriented according to the alignment with the chromosome-level GRC assembly of *L. malacca*, thus maximizing collinearity between the species. This figure thus represents a lower estimate of the amount of rearrangements on GRC. The color of the connection is determined by the reference position from left to right.

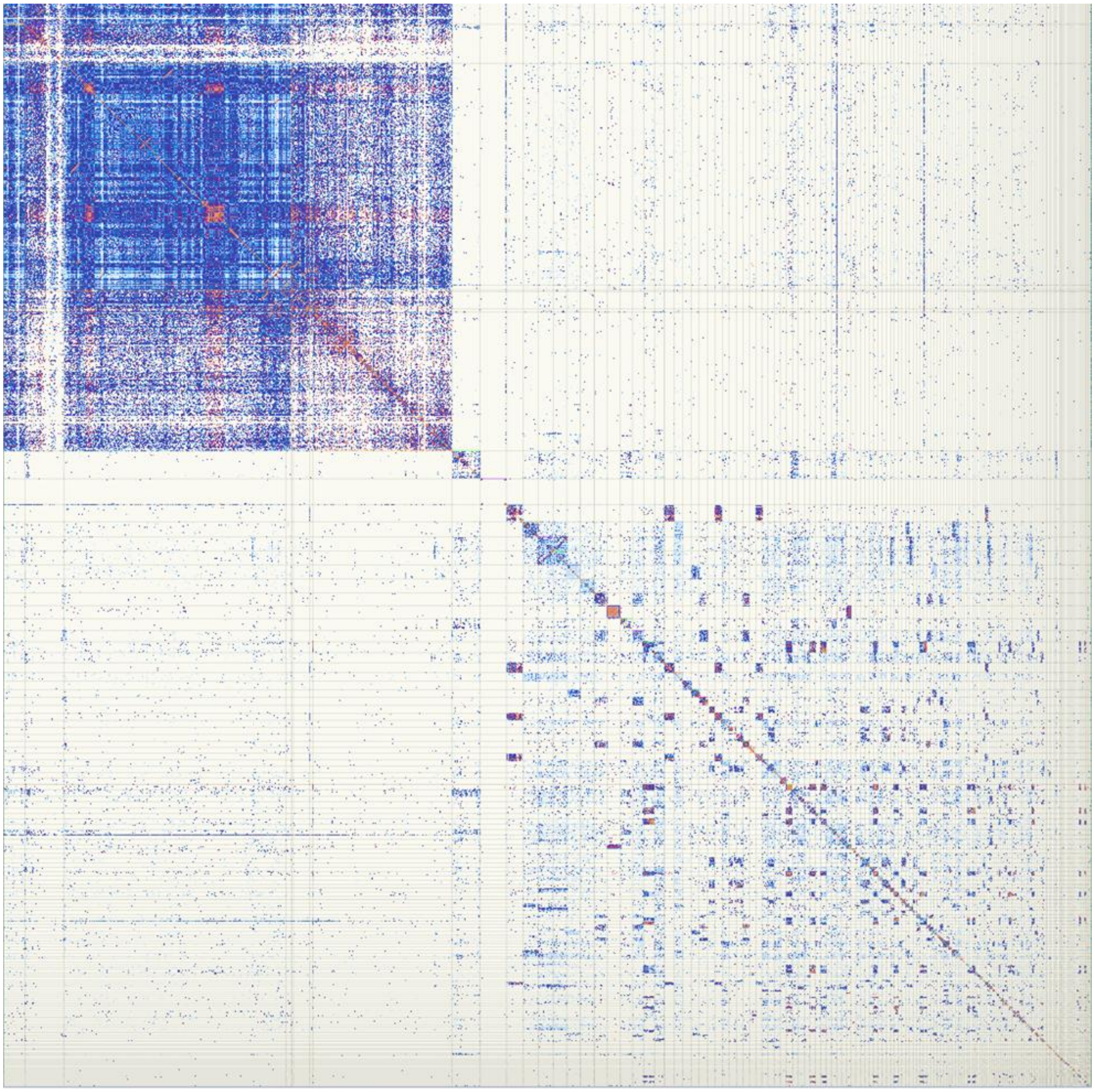

**Supplementary Figure 12: Omni-C map of connections between putative *L. malacca* GRC contigs.** Contigs finally identified as being part of the GRC are on the left and have high inter contig alignments rates from the Omni-C data.

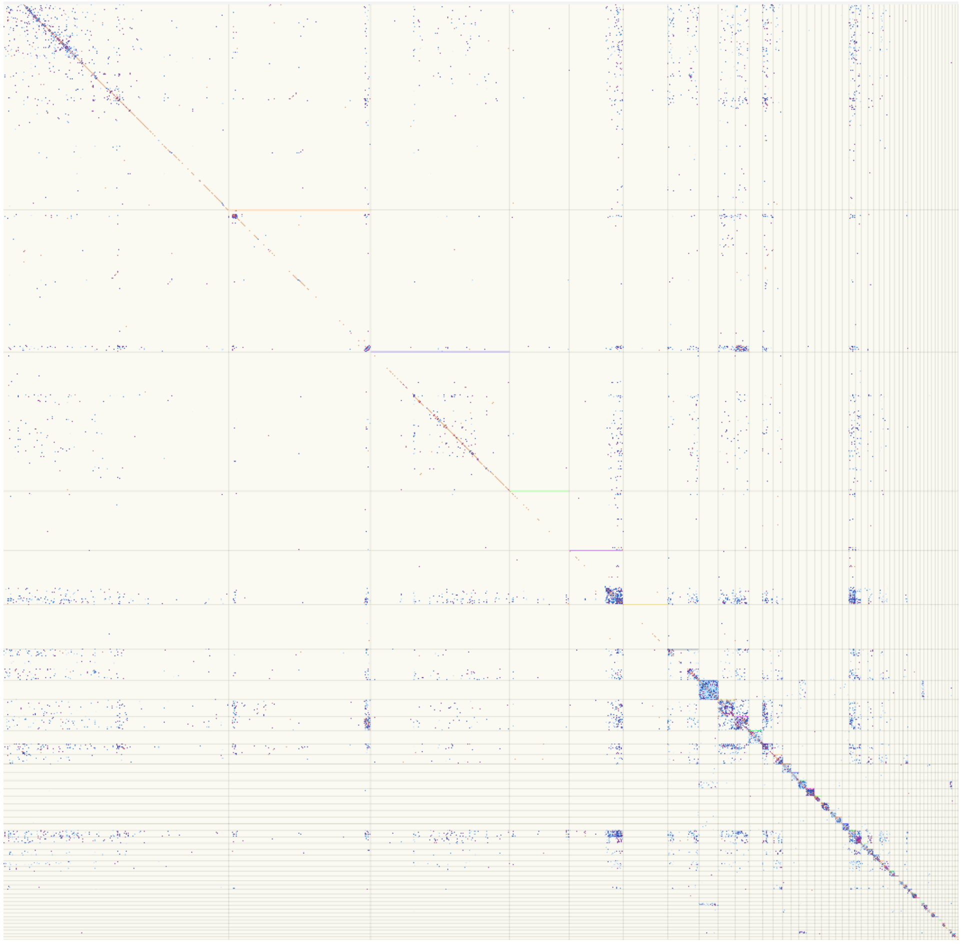

**Supplementary Figure 13: Omni-C map of connections between putative *L. punctulata* GRC contigs.** The sequencing depth and unique mapping rate eventually resulted in a low signal strength.
